## Supplement for "Hierarchical progressive learning of cell identities in single-cell data"

**Supplementary Note 1**

When matching the cell populations from two datasets, we distinguish five options: simple, multiple columns, multiple rows, complex, and impossible. When describing the different scenarios within these options, we sometimes make a distinction between leaf nodes and internal nodes. Here, it is important to remember that only *T1* can have internal nodes since this is the tree that is updated. *T2* is always a flat classification tree, so only consists of the root node and leaf nodes.

*Simple*

In this scenario, we find a unique match between a cell population, $P_i$, from dataset 1 and a cell population, $P_j$, from dataset 2. As as consequence, $X_{j,I}$ will be 1 or 2 and the rest of row *j* and column *i* in *X* are zero. Within this scenario, there are three different options:

1. Both cell populations are leaf or internal nodes. This indicates a perfect match. The tree is not updated, but the labels of $P_j$ are renamed to $P_i$ (Figure S4A). This is the same scenario as the 'perfect match' scenario described in the main text.
2. $P_i$ is a leaf or internal node, but $P_j$ is the root node of *T2*. This indicates that $P_i$ is missing in dataset 2. The node, however, is already in the tree, so it is not updated (Figure S4B).
3. $P_i$ is the root of *T1*, but the $P_j$ is a leaf node. This indicates that $P_j$ is missing in dataset 1. The cell population is thus also not in the tree yet, so we will add it as a child to the root (Figure S4C). This is the same scenario as the 'new population' scenario described in the main text.

*Multiple rows*

In this scenario, a cell population, $P_i$, from dataset 1 matches multiple populations from dataset 2. In *X* there will be multiple non-zero values in column *i.* Here, we distinguish two different scenarios:

1. $P_i$ matches only cell populations from dataset 2 that are leaf node. We consider the cell populations from dataset 2 subpopulations of $P_i$, so we add them as descendants to $P_i$ (Figure S5A). This is the same scenario as the 'splitting nodes' scenario described in the main text.
2. The root node of *T2* is also involved. We simple ignore this node and for the rest do the same as above (Figure S5B-C).

*Multiple columns*

This scenario is quite similar to the multiple rows scenario. Here, however, multiples populations from dataset 1 match one cell population, $P_j$, of dataset 2. In *X* there will be multiple non-zero values in row *j.* This scenario is a little more complex since the populations from dataset 1 does not have to be leaf nodes or the root node, but there can also be internal nodes in this tree. Here, we distinguish three different scenarios:

1. The root node of *T1* and *T2* are not involved, so multiple cell populations, which can be leaf or internal nodes, from dataset 1 match $P_j$. We consider the cell populations from dataset 1 subpopulations of $P_j$, so we need to add $P_j$ as a parent node to these cell populations. (Figure S6A). This is same scenario as the 'merging nodes' scenario described in the main text. It could be, however, that this node already exists in this tree. (Figure S6B). If this is the case, we have a perfect match between a node from tree 1 and tree 2, so we do not have to update the tree, but we only have to update the labels of $P_j$.
2. Besides leaf or internal nodes, the root of *T1* is involved. This indicates that $P_j$ is 'bigger' than the cell populations from dataset 1 as part of it is unlabeled. Therefore, we add $P_j$ as a

descendant to the root of *T1*. Next, we rewire the involved cell populations from dataset 1 such that they become descendants of $P_j$ (Figure S6C).

3. The root node of *T2* is involved. This indicates that multiple cell populations from dataset 1 are missing in dataset 2. These nodes, however, are already in the tree, so the tree can remain the same (Figure S6D).

*Complex*

The scenarios described above were all relatively easy. A cell population from one dataset matches either one or multiple cell populations from another. It could also happen, however, that multiple cell populations from dataset 1 match multiple cell populations from dataset 2 (Figure S7). As a consequence, there will a certain place $X_{j,I}$ which is either 1 or 2 and there are two or more non-zero values in the corresponding row *j* and column *i.* Here, we distinguish three different scenarios:

1. The root node of *T1* is involved. We just assume that the boundary should be adjusted and this is automatically done, so we remove this `1' from the table (Figure S7A). If the situation is still complex after the one is removed, we continue to scenario 2 or 3. If not, we treat it as a multiple rows problem as explained above.
2. The root node of *T2* is involved. Again, we just assume that the boundary should be adjusted, so we remove this `1' from the table (Figure S7B). If the situation is still complex after the one is removed, we continue to scenario 3. If not, we treat it as a multiple columns problem as explained above.
3. Multiple leaf/internal nodes of dataset 1 are involved and multiple leaf nodes of dataset 2. We can only solve this if the 'complex' cell population, $P_i$, of dataset 1 is not a leaf node. Otherwise we are dealing with an impossible scenario which is described below. If the complex node is an internal node, we attach the involved cell populations of dataset 2 as descendants to the complex node (splitting scenario) and attach the involved cell populations of dataset 1, except for $P_i$, to $P_j$ (Figure S7C).

*Impossible*

Sometimes, it could be impossible to match the labels from two datasets. Something could have gone wrong during the clustering, e.g. a population 1 and 2 from dataset 1 match population A from dataset 2, but population 2 also matches population C from dataset 2 (Figure S8A). Here, population A and C should be merged into population 2, but population A should also be split into population 1 and 2. Population 2, however, cannot be added to the tree twice.

It could also be that dataset 2 contains labels at a different resolution, e.g. that population B is a subpopulation of population A (Figure S8B). This is not what we assumed and thus impossible to match.

Both scenarios occur when a leaf node from dataset 1 is at a crossing of multiple rows and multiple columns (i.e. a complex situation). An extra difficulty is that there are thus multiple situations that could explain this. All of these situation are not what we desired and thus we call it impossible and do nothing.

**Supplementary Note 2**

If there is a complex scenario that cannot be solved immediately, matrix *X* will be changed into a strict matrix. In the strict matrix, only reciprocal matches are considered, so all ‘1’s’ are turned into ‘0’. There are some exceptions to this rule.

- A population can never have a reciprocal match with the root, so these ‘1’s’ are never removed.
- If a population from a dataset has only one match, it is also never removed. Consider the following example: If population P1 of Dataset 1 is only predicted to be Population Q of Dataset 2, we know that P1 should be a match with Q as it cannot be matched with any other population or with the root. It could be that this match is not reciprocal if population Q has many different subpopulations (e.g. P1, P2, P3, P4). Imagine that population P2 is really big. Almost all cells of population Q will be predicted to be P2 and so the matches with P1 (and P3 and P4) are missed because of the matching threshold. In case there is a complex scenario caused by any other population (maybe P2 or P3 or P4), we still know that P1 is a subpopulation of Q, since that was super clear and didn’t cause any complexity.

**Supplementary Note 3**

Current scRNA-seq data simulators cannot simulate hierarchical data, so we simulated this dataset step by step (Figure S1B).

First, we simulated the expression of 3,000 genes for 9,000 cells. For this simulation, the cells were divided into three groups. The 3,000 simulated genes represent genes that are differentially expressed between the cell populations at a low resolution, so for example B cells vs. T cells. Next, we simulated *another* 3,000 genes for the *same* 9,000 cells. Now, the cells were divided into five groups. Here, the differentially expressed genes represent genes that distinguish cell populations at a slightly higher resolution, so for example CD4+ T cells vs. CD8+ T cells. We repeated this step for another set of 3,000 genes, but now there were six populations. The third dataset represents the highest resolution, so for instance CD4+ memory T cells vs. CD4+ naïve T cells.

Together this resulted in a dataset of 9,000 cells and 9,000 genes. The cells were labeled at three resolutions. There was some inconsistency between the labels at the different resolutions (e.g. some cells were labeled as ‘Group12’, ‘Group3’, ‘Group3’). We removed these cells from the dataset, which resulted in a final dataset of 8,839 cells and 9,000 genes.

**Supplementary Figures**

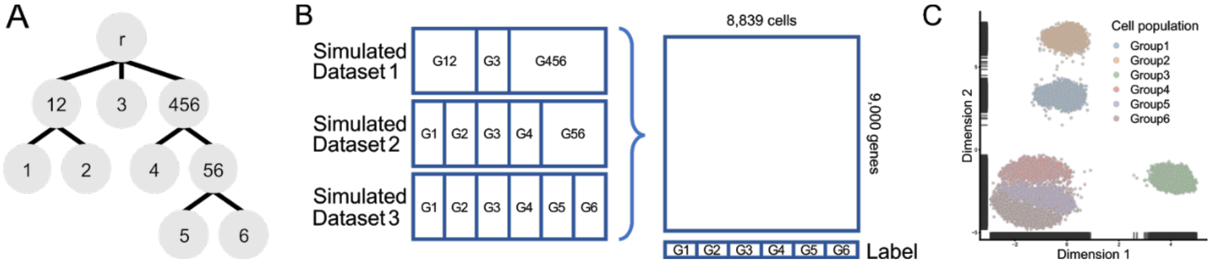


**Figure S1** (A) Classification tree for the simulated dataset. (B) We simulated three datasets separately and concatenated them in one dataset. The labels and their proportion are indicated in the simulated datasets. (C) UMAP of the final dataset.

A

B

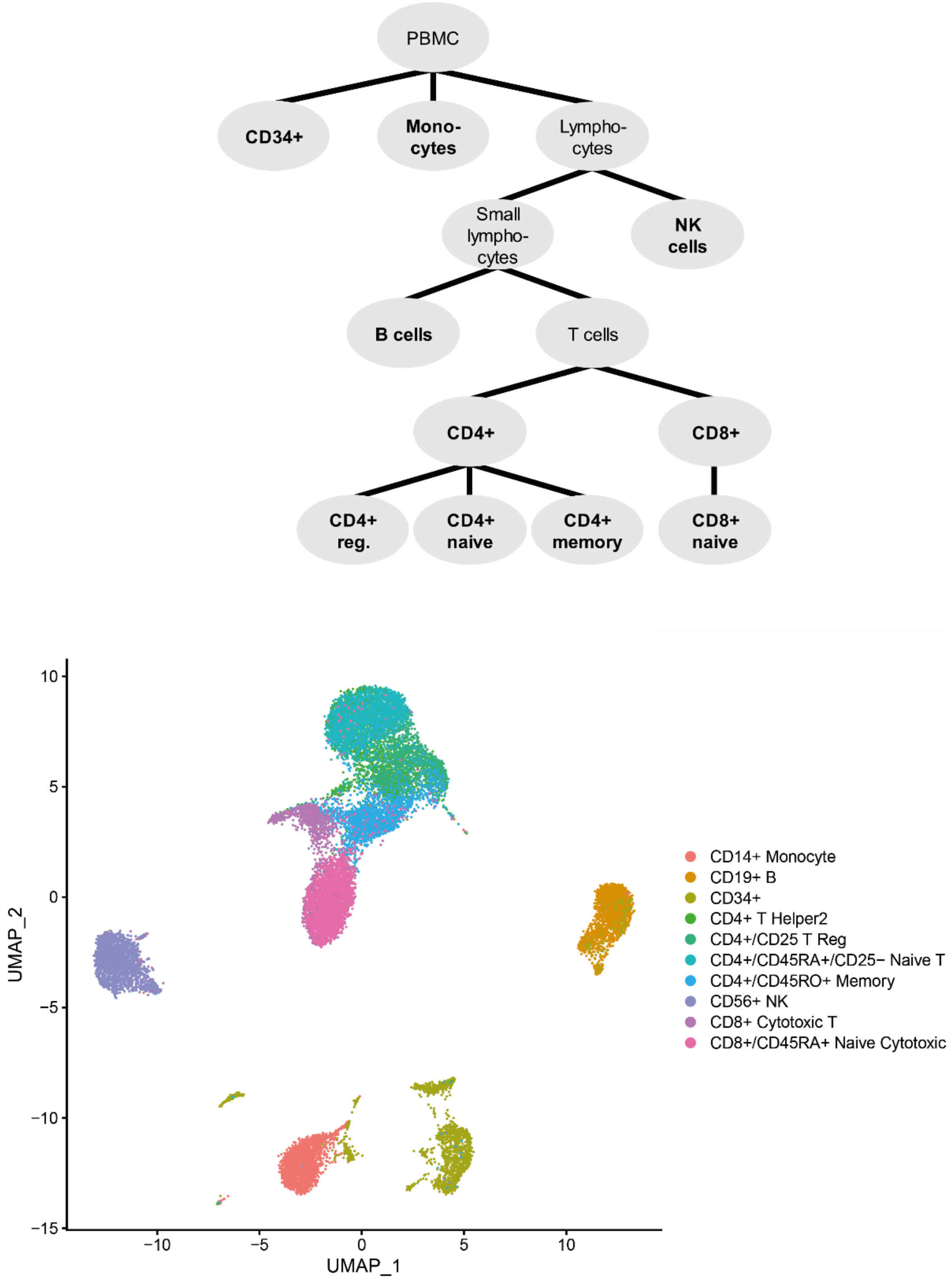


**Figure S2** (A) Classification tree for the PBMC-FACS dataset. Bold names indicate cell populations that exist in our dataset. (B) UMAP embedding of the PBMC-FACS data

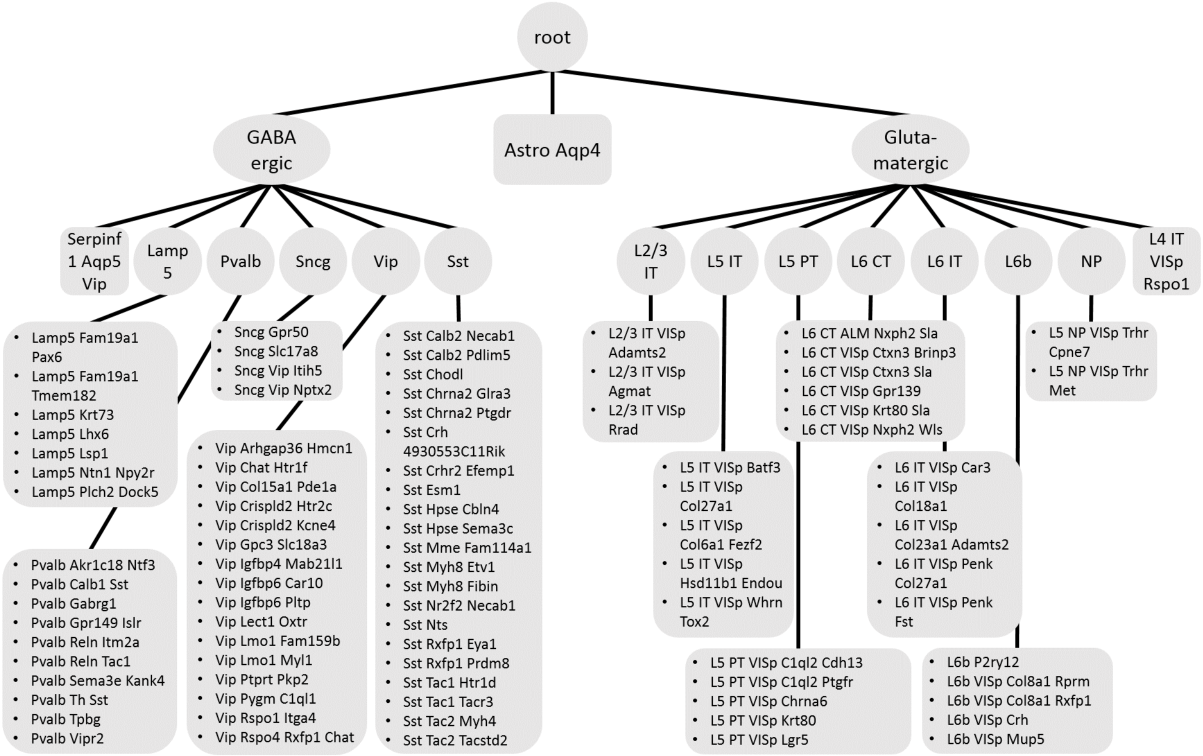


**Figure S3** Classification tree of the AMB dataset. The circular and rectangle nodes indicate internal and leaf nodes respectively. If multiple leaf nodes are descendants of the same internal node, they are placed in the same rectangle.

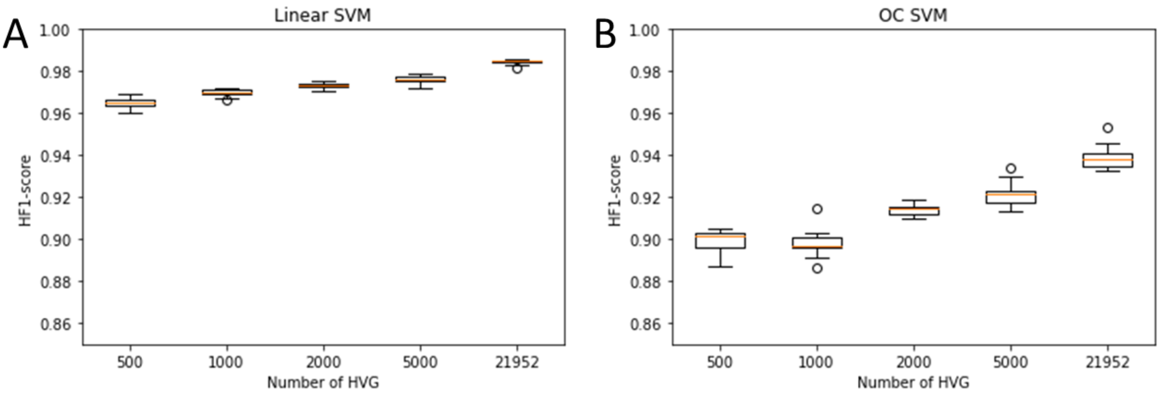


**Figure S4** Effect of selection HVG on the classification performance on the PBMC-FACS dataset when using the (A) linear SVM and (B) one-class SVM.

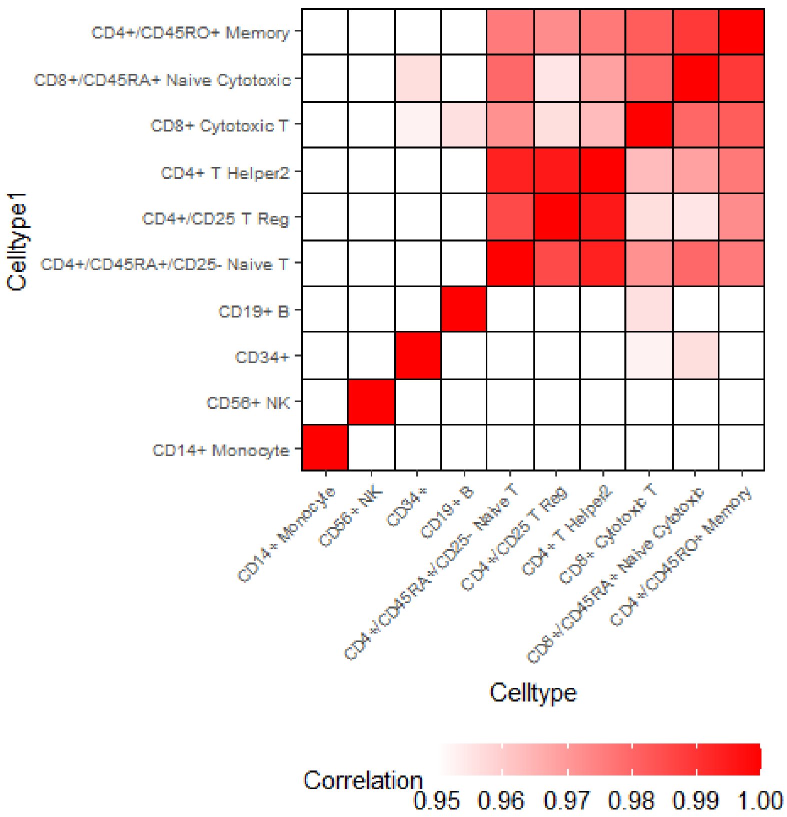


**Figure S5** Pearson correlation between the cell population in the PBMC-FACS dataset.

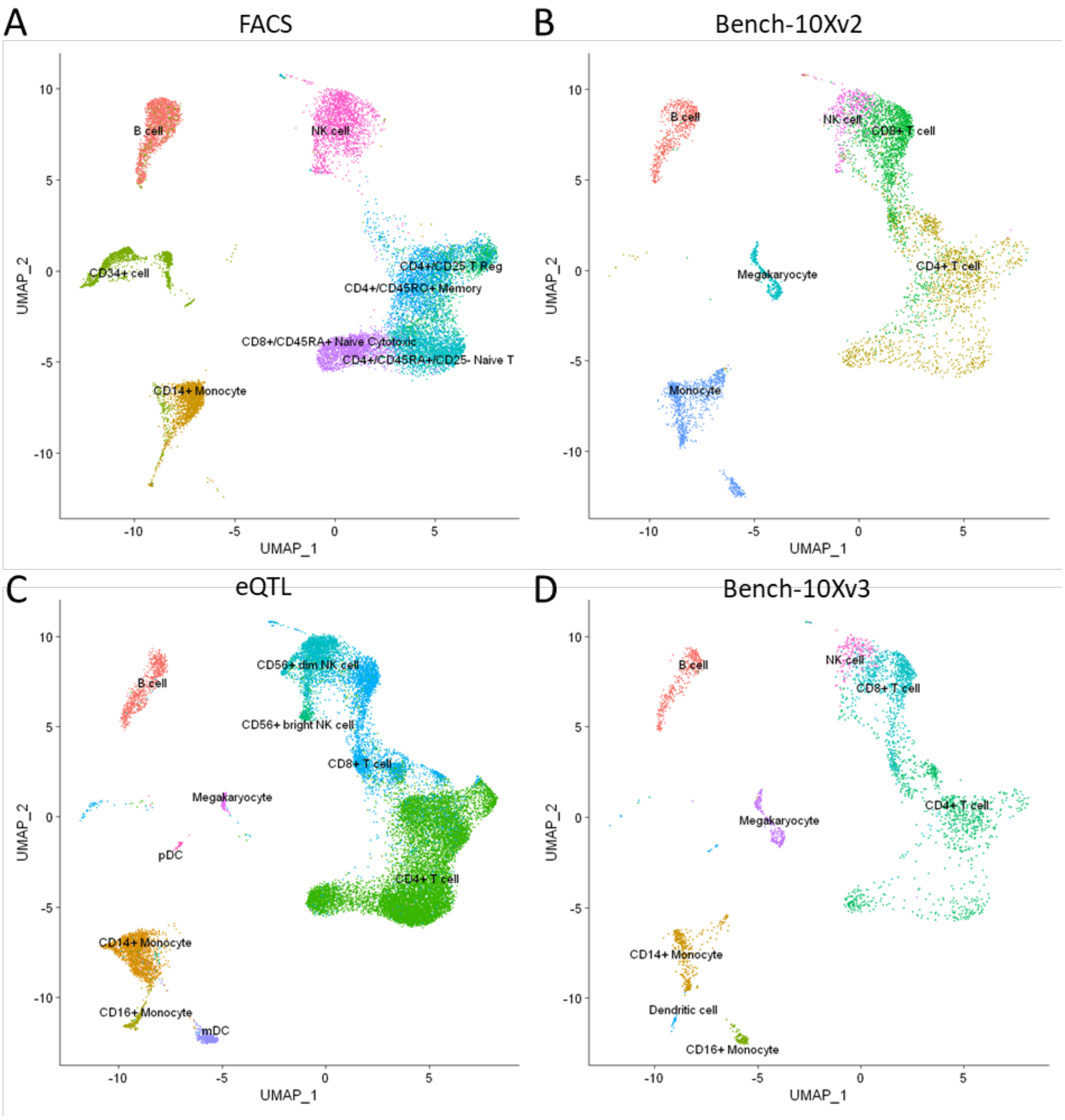


**Figure S6** UMAPs showing the cell populations of the (A) FACS, (B) Bench-10Xv2, (C) eQTL, (D) Bench-10Xv3 datasets after integration with Seurat.

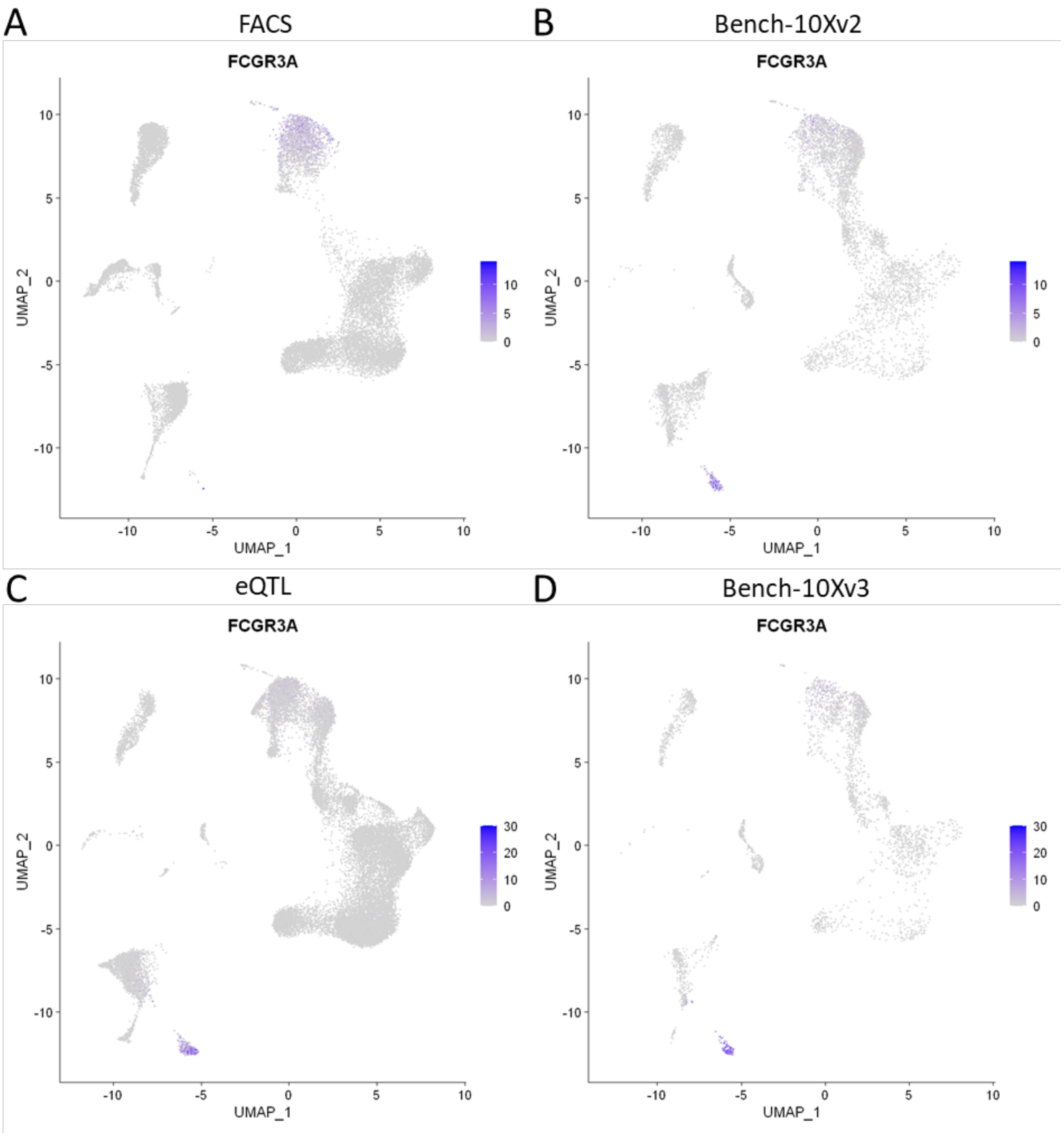


**Figure S7** UMAPs showing the expression of *FCGR3A* in the PBMC datasets. We visualized the original counts of *FCGR3A,* so before integration. *FCGR3A* is a marker gene for CD16+ monocytes. In the eQTL dataset, *FCGR3A* is expressed in the cells labeled as mDC (Figure S7).

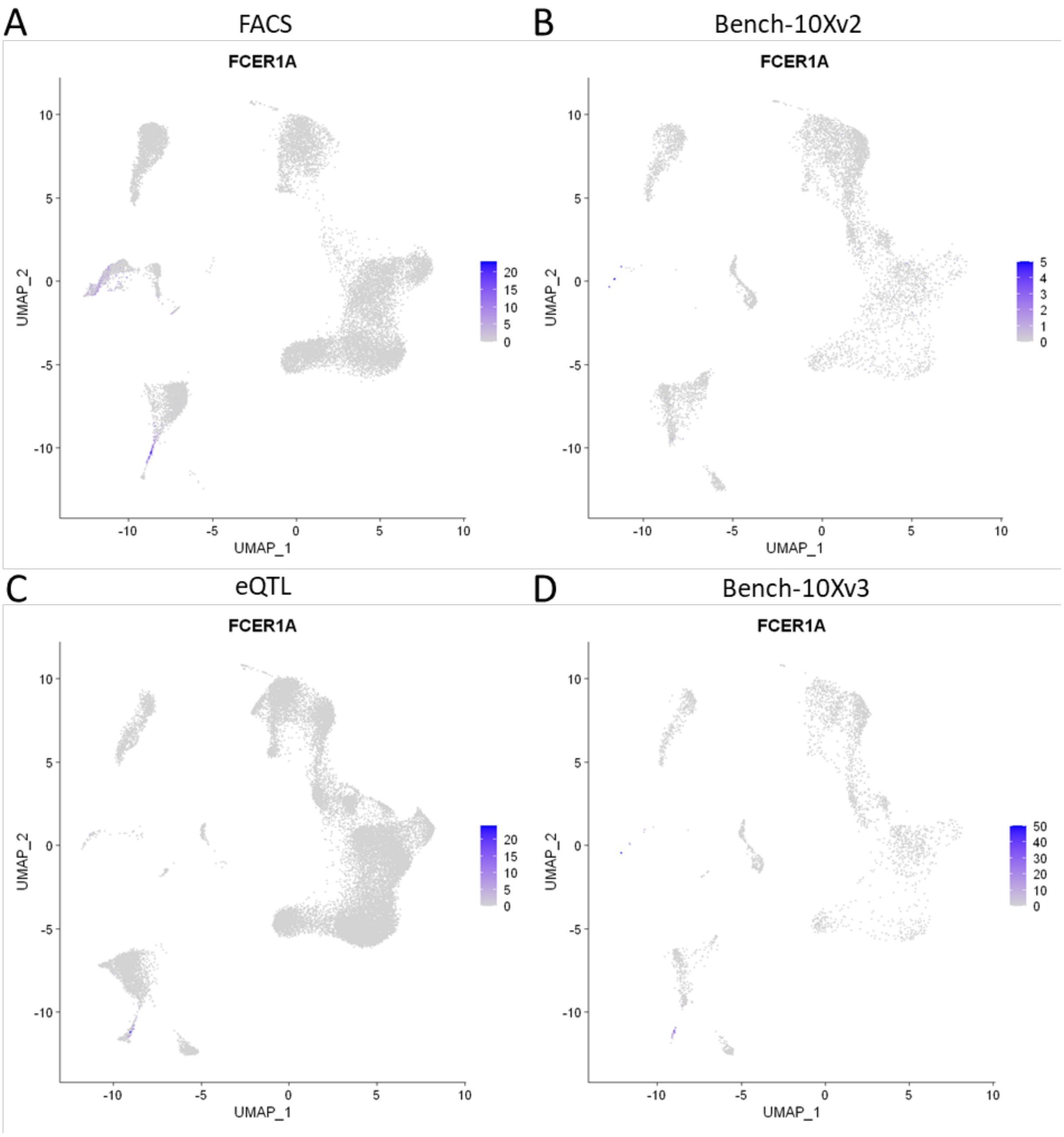


**Figure S8** UMAPs showing the expression of *FCER1A* in the PBMC datasets. We visualized the original counts of *FCER1A,* so before integration. *FCER1A* is a marker gene for mDC. In the eQTL dataset, *FCER1A* is lowly expressed in the cells labeled as CD16+ monocytes (Figure S7). In the FACS dataset, part of the cells labeled as CD14+ monocytes also expressed *FCER1A.*

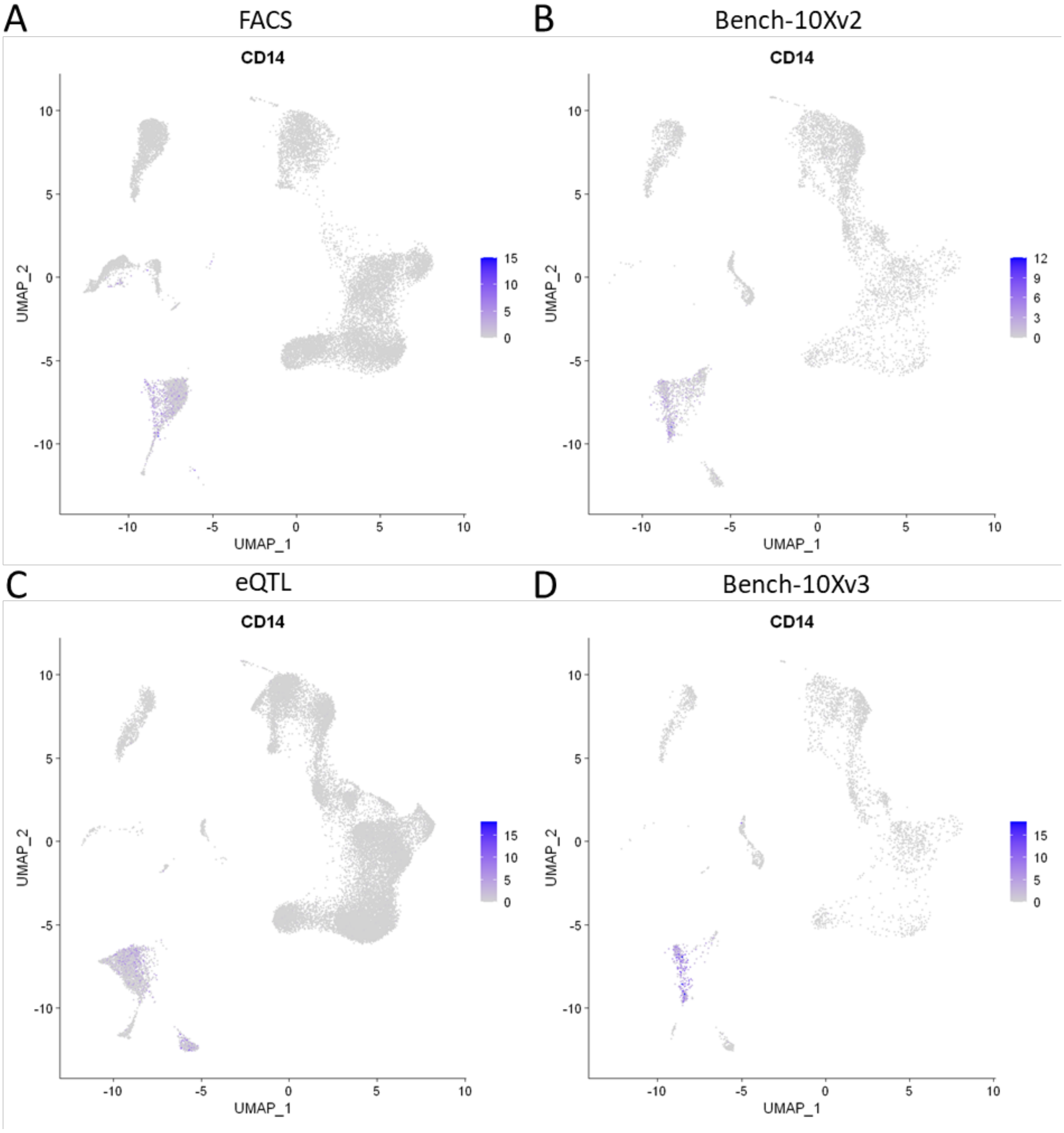


**Figure S9** UMAPs showing the expression of *CD14* in the PBMC datasets. We visualized the original counts of *CD14,* so before integration. *CD14* is a marker gene for CD14+ monocytes. In the FACS dataset, part of the cells labeled as CD14+ monocytes express no CD14+ (Figure S7).

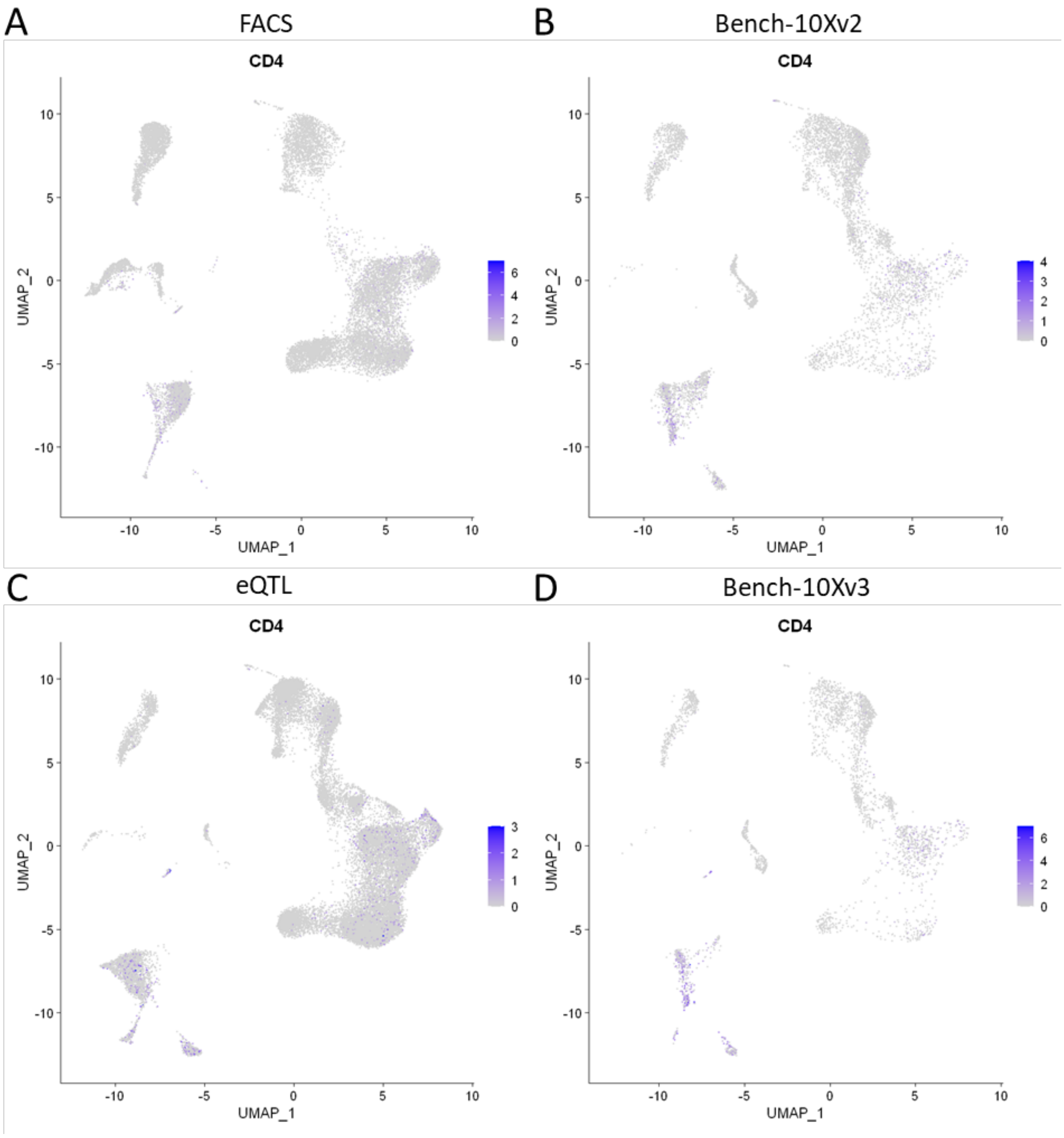


**Figure S10** UMAPs showing the expression of *CD4* in the PBMC datasets. We visualized the original counts of *CD4,* so before integration. *CD4* is a marker gene for CD4+ T-cells and monocytes. In the eQTL and Bench-10Xv2 dataset, *CD4* is only lowly expressed in the CD4+ T-cells at the right of the cluster and expression is missing at the bottom-left part of the cluster (Figure S7).

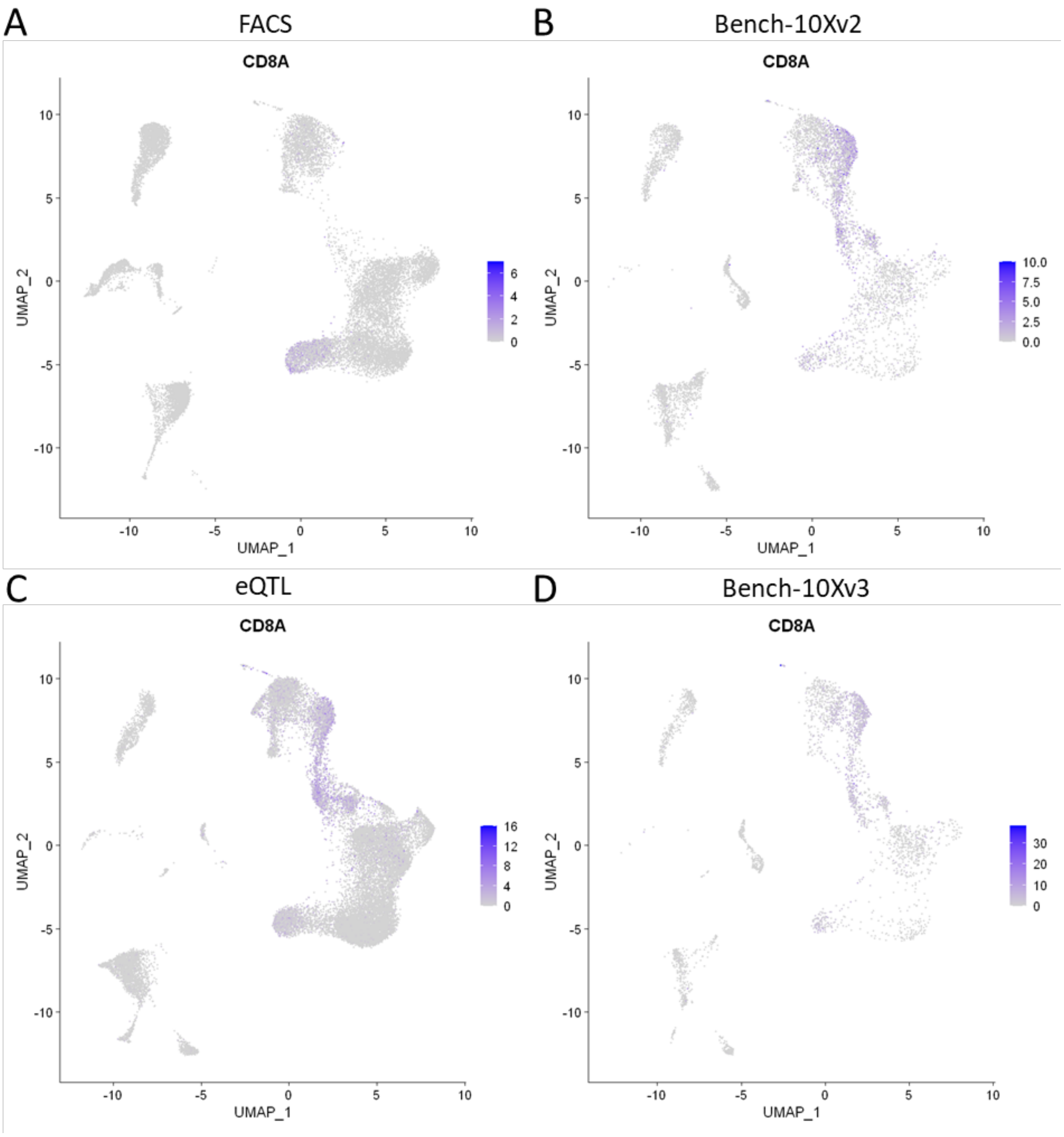


**Figure S11** UMAPs showing the expression of *CD8A* in the PBMC datasets. We visualized the original counts of *CD8A,* so before integration. *CD8A* is a marker gene for CD8+ T-cells. In the eQTL and Bench-10Xv2 dataset, *CD8A* is also expressed in the CD4+ T-cells at the bottom-left part of the cluster (Figure S7).

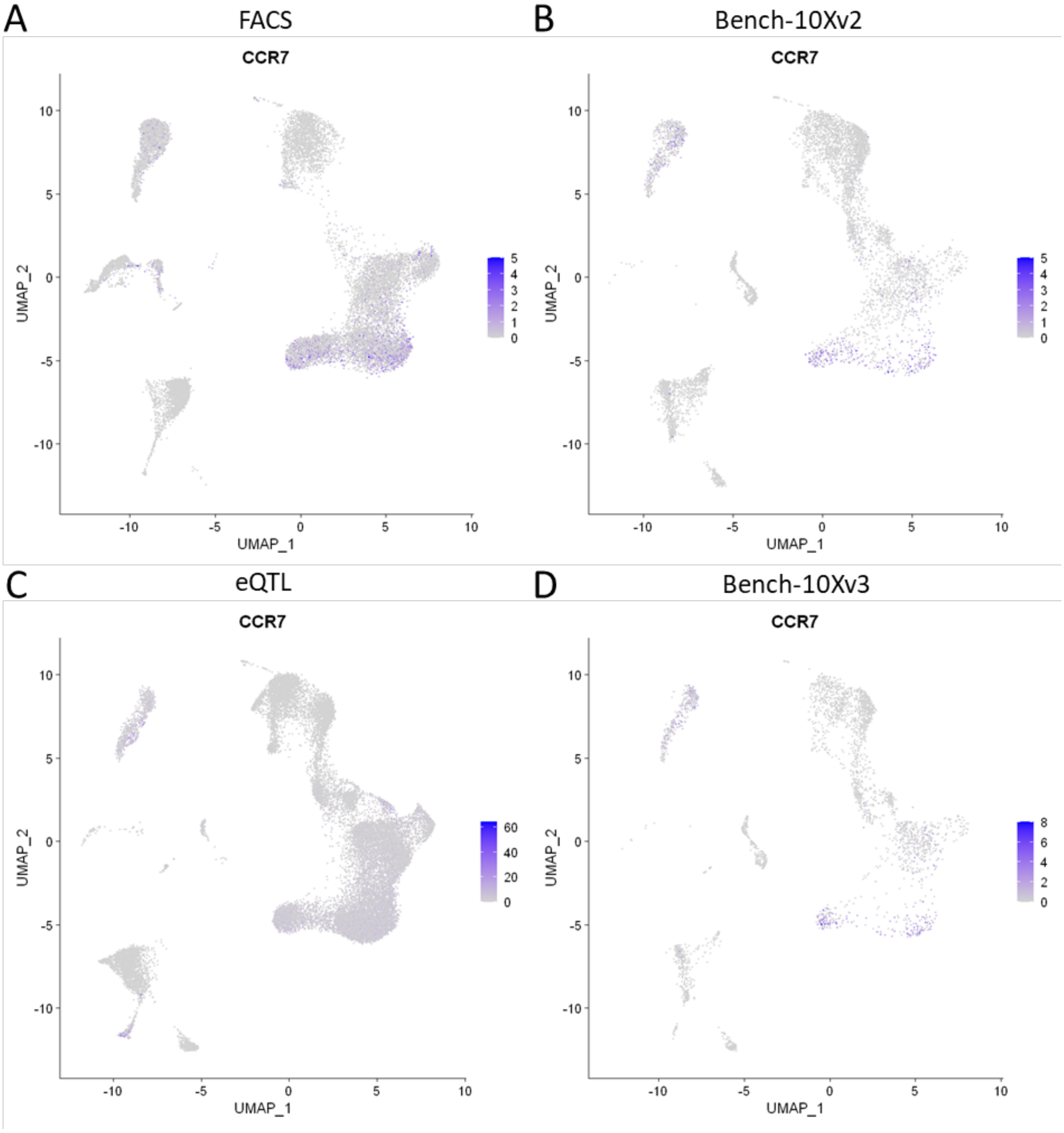


**Figure S12** UMAPs showing the expression of *CCR7* in the PBMC datasets. We visualized the original counts of *CCR7,* so before integration. *CCR7* is a marker gene for naive T-cells.

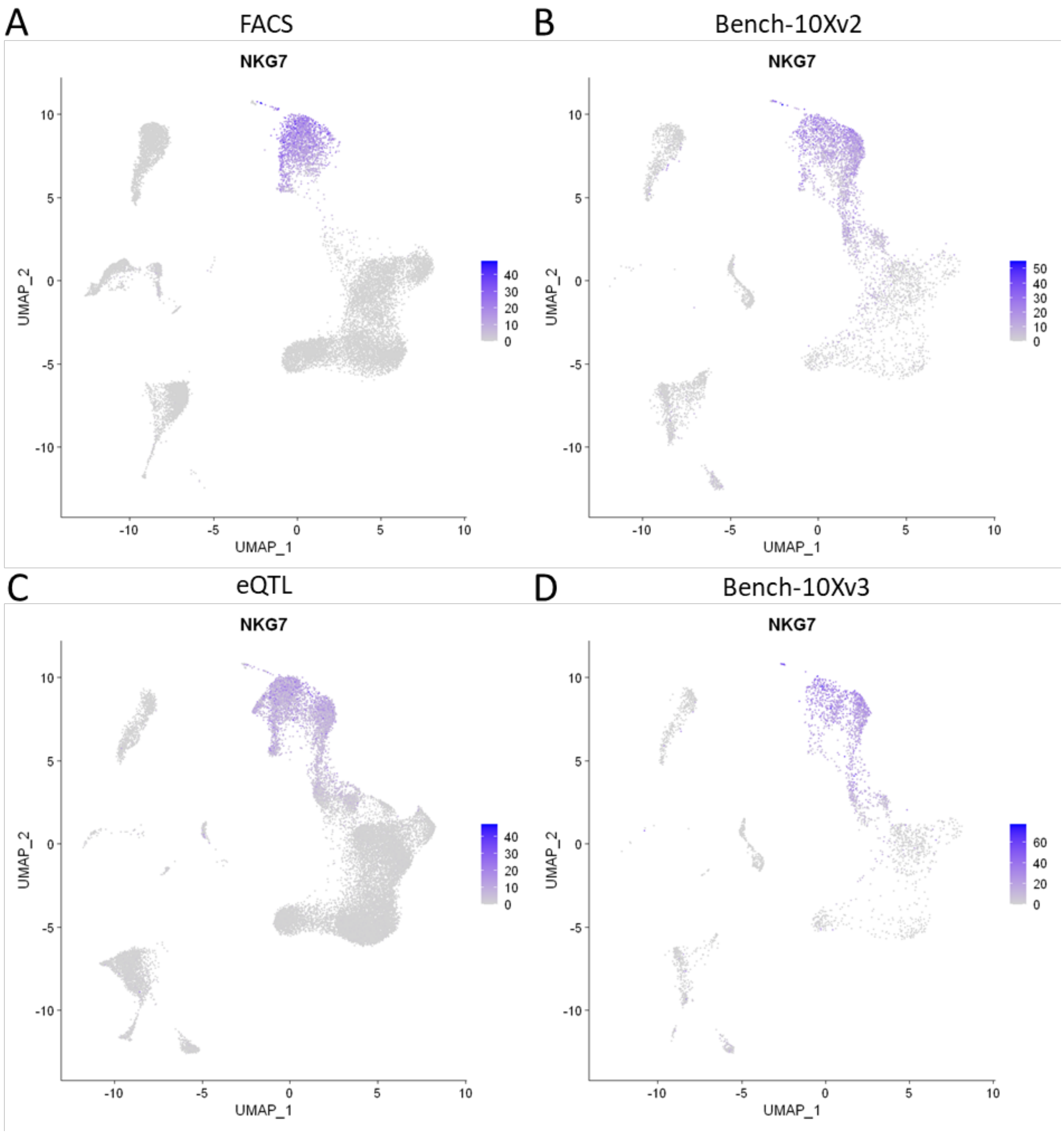


**Figure S13** UMAPs showing the expression of *NKG7* in the PBMC datasets. We visualized the original counts of *NKG7,* so before integration. *NKG7* is a marker gene for NK cells and cytotoxic T-cells.

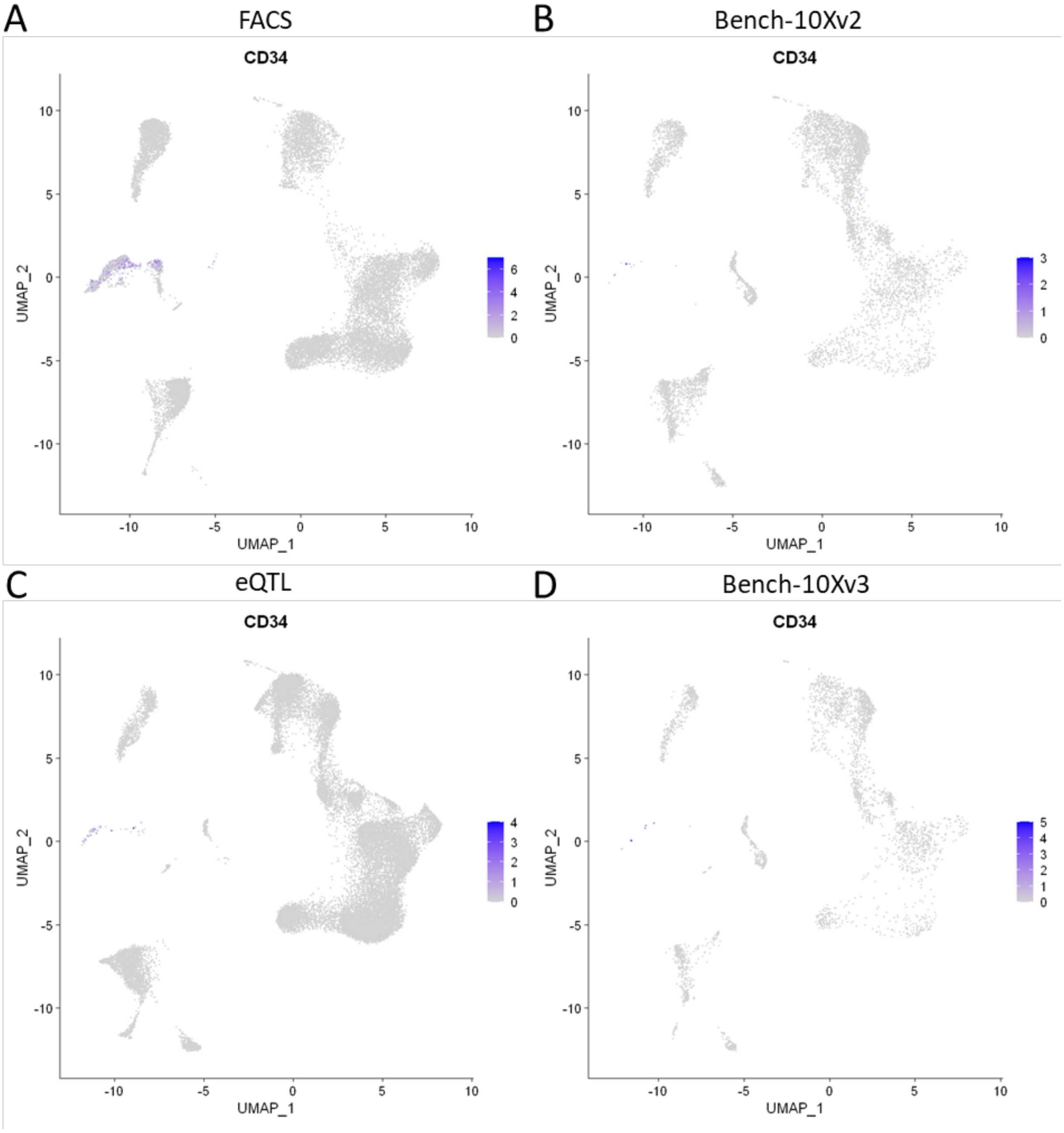


**Figure S14** UMAPs showing the expression of *CD34* in the PBMC datasets. We visualized the original counts of *CD34,* so before integration. *CD34* is a marker gene for CD34+ cells.

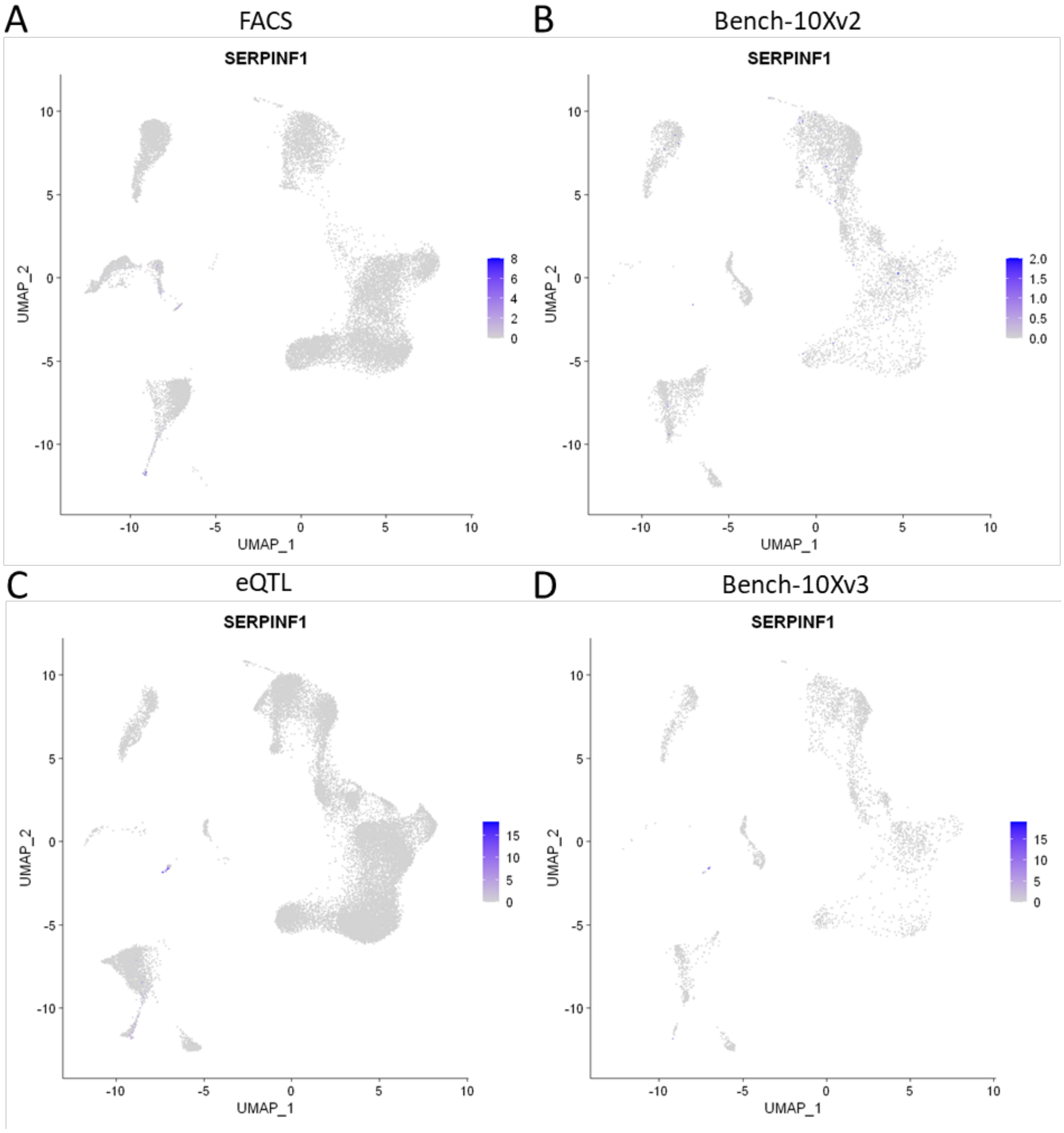


**Figure S15** UMAPs showing the expression of *SERPINF1* in the PBMC datasets. We visualized the original counts of *SERPINF1,* so before integration. *SERPINF1* is a marker gene for pDC.

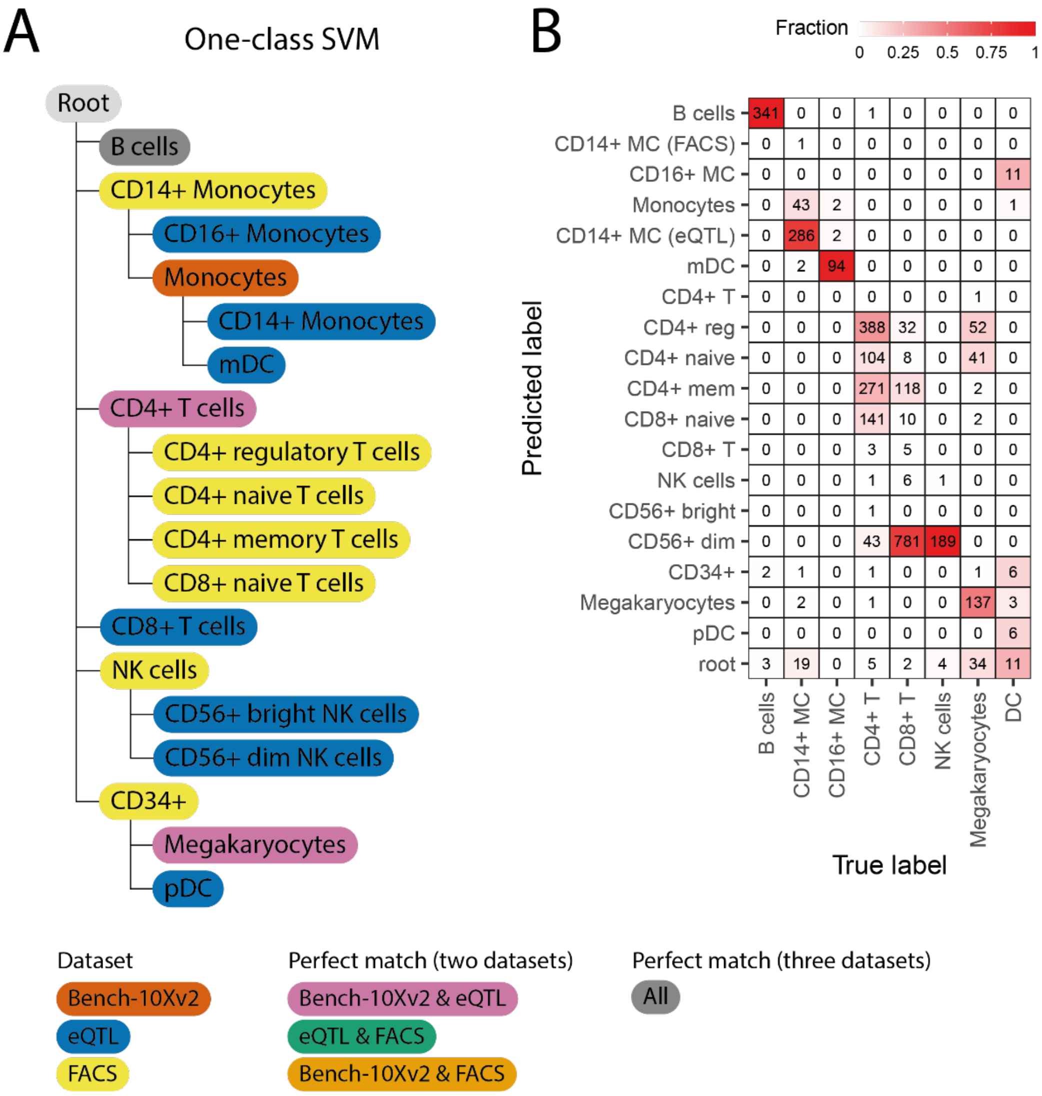


**Figure S16** (A) Constructed classification tree when using a one-class SVM during the PBMC inter-dataset experiment. The color of a node represents the dataset(s) of the cell population. If a color refers to multiple datasets, this indicates that the populations from these datasets had a perfect match. The NK and CD8+ T-cell populations from the PBMC-Bench10Xv2 are missing from the tree. (B) Confusion matrix when using the constructed classification tree to predict the labels of PBMC-Bench10Xv3.

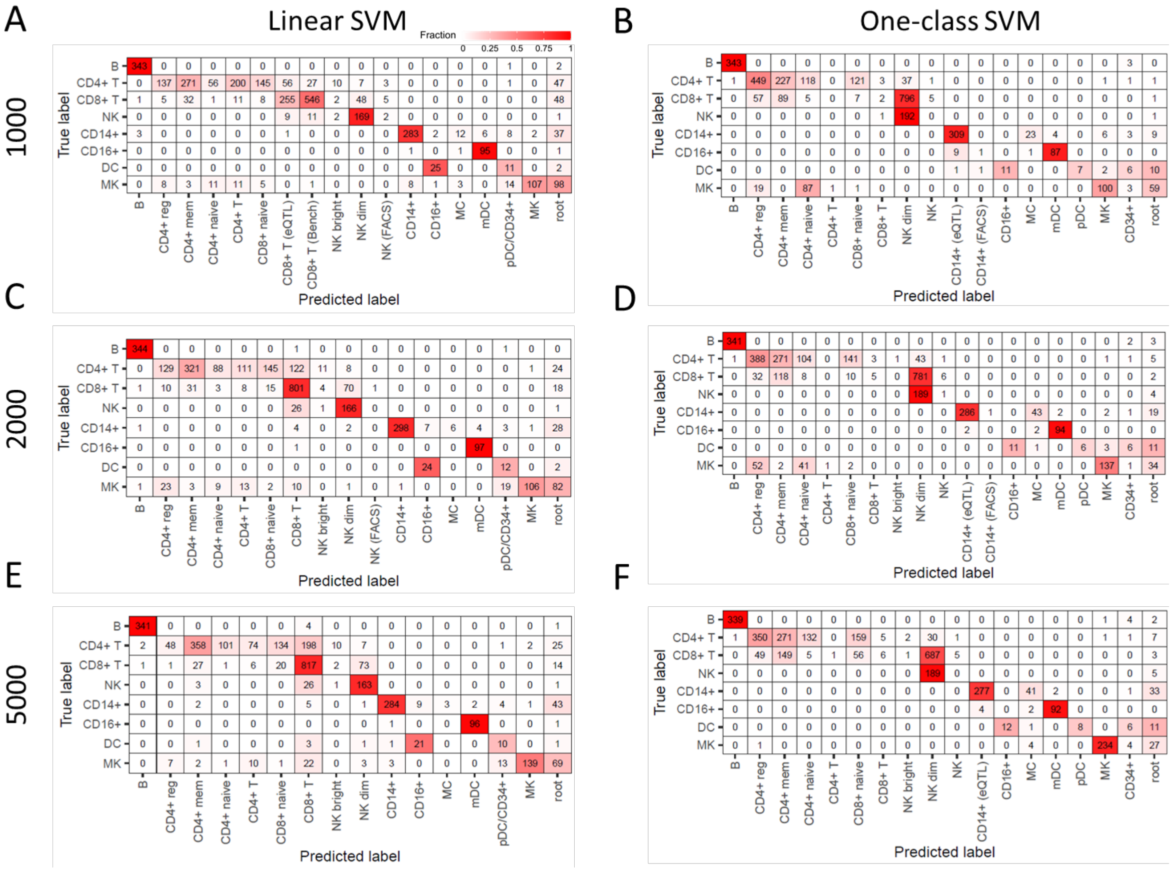


**Figure S17** Confusion matrix when using the constructed classification tree to predict the labels of PBMC-Bench10Xv3 with linear and one-class SVM (columns) when varying the number of anchors to integrate the dataset (rows).

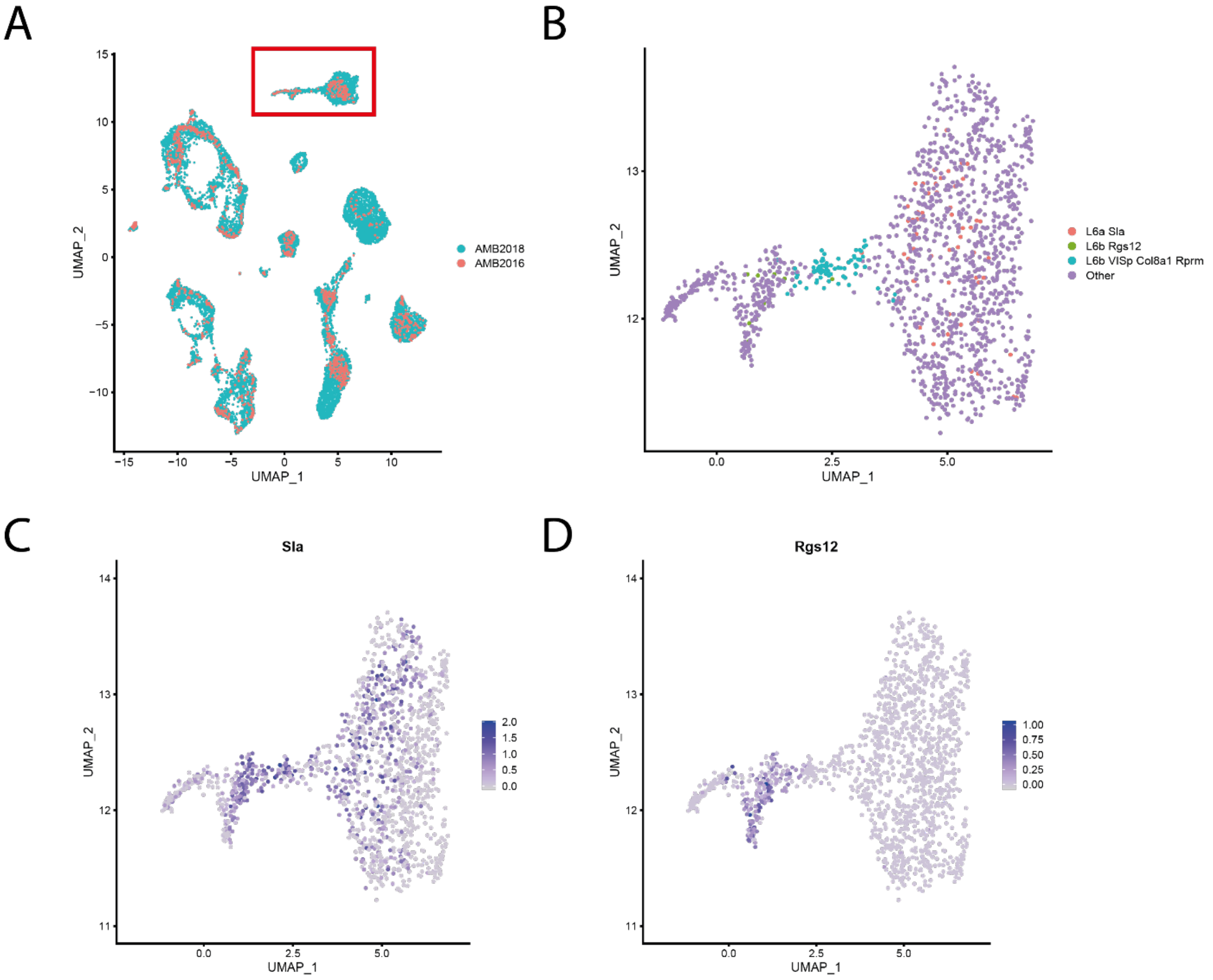


**Figure S18** (A) UMAP embedding showing the AMB2016 and AMB2018 datasets after data integration. The red rectangle shows the cells we zoomed in on to visualize the expression of marker genes. (B) Different cell populations within this part of the UMAP. (C-D) Expression of *Sla* and *Rgs12.* The 'L6b VISp Col8a1 Rprm' population does not express *Rgs12,* but does express *Sla,* which support the match with 'L6a Sla'  instead of 'L6b Rgs12'.

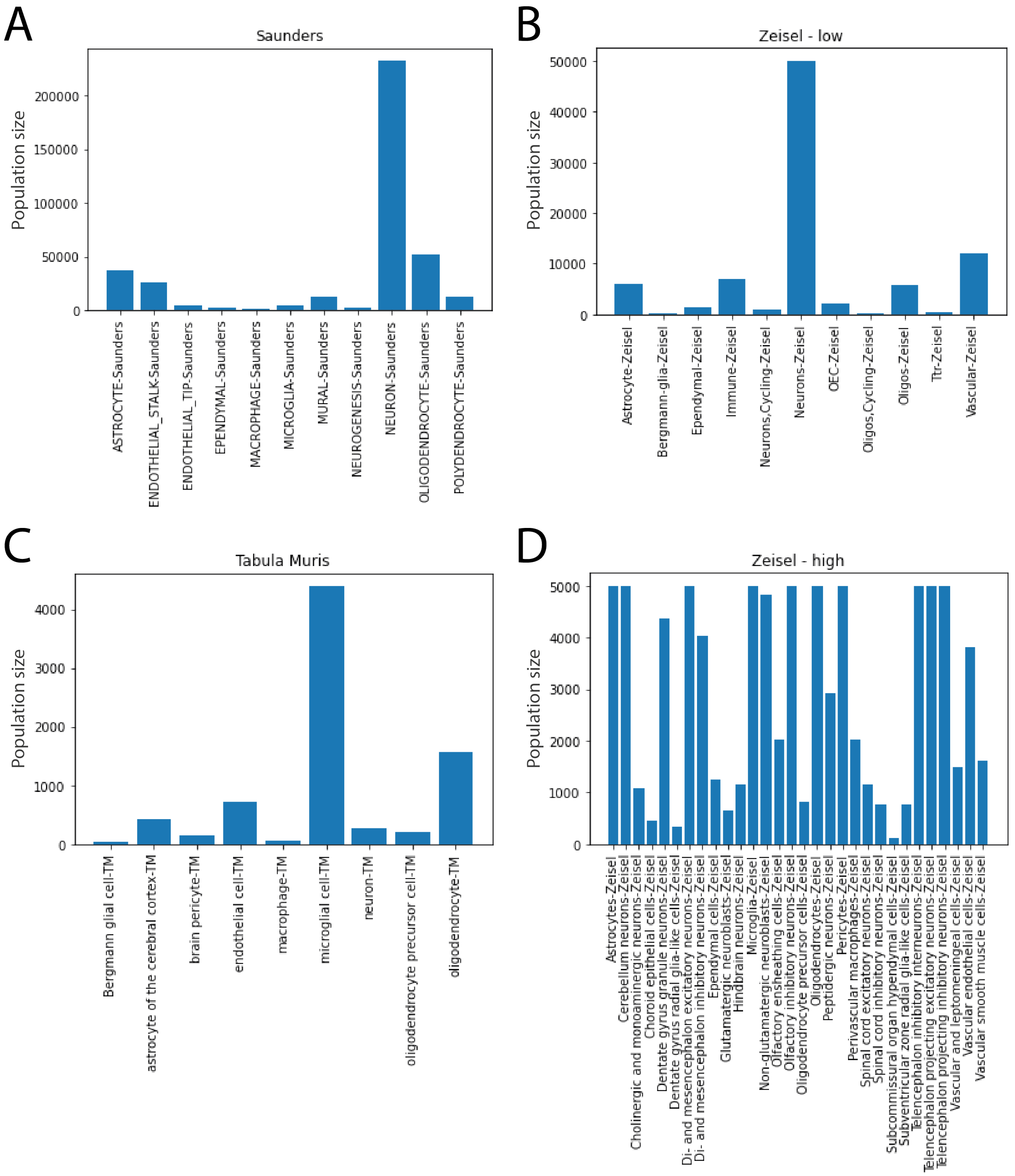


**Figure S19** Size of the cell populations in the (A) Saunders, (B) Zeisel – low resolution, (C) Tabula Muris, and (D) Zeisel – high resolution dataset. When comparing the size of the difference populations, notice the different scales on the y-axis.

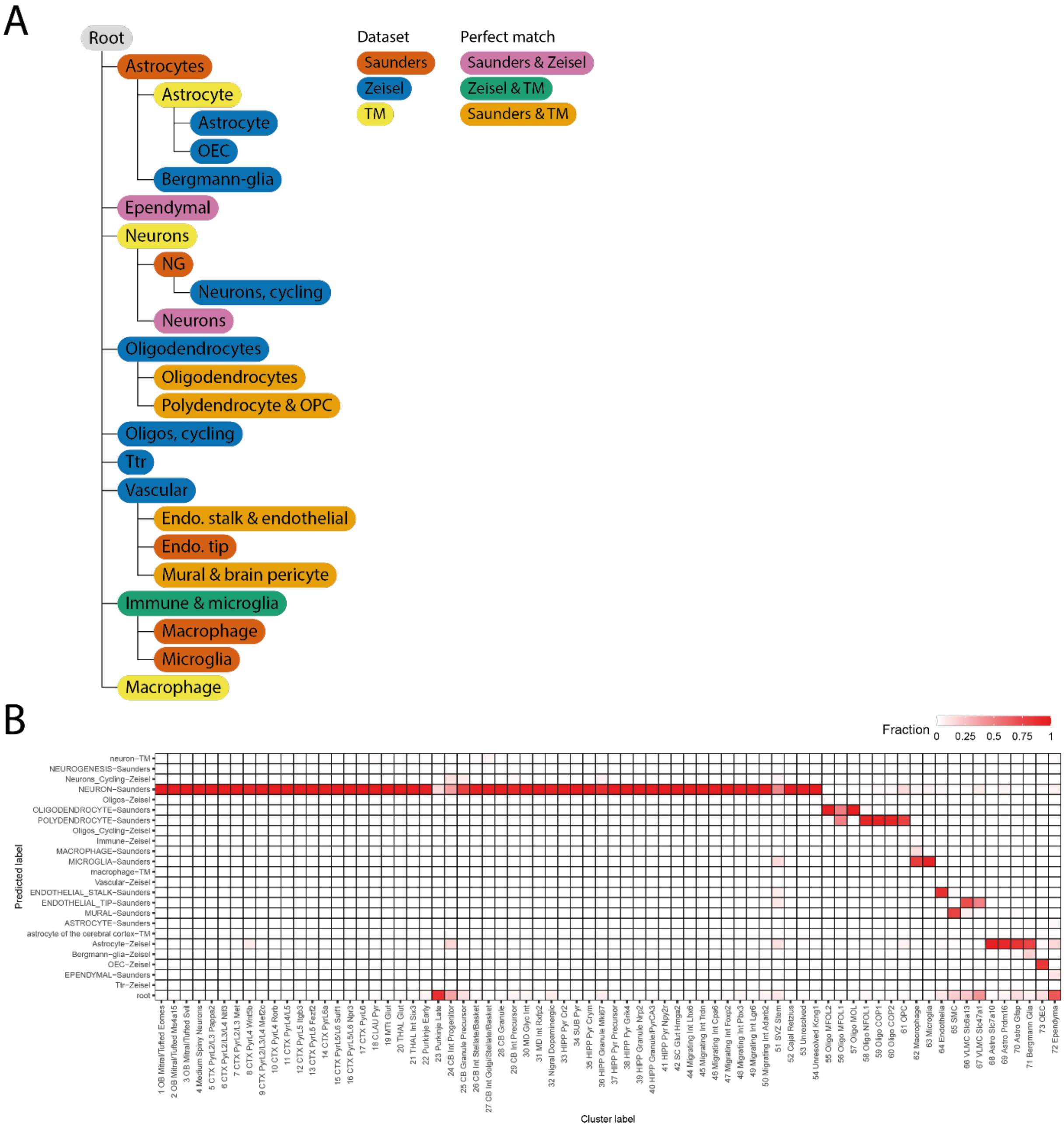


**Figure S20** (A) Learned hierarchy when applying *scHPL* with a one-class SVM to the Saunders, Zeisel (low resolution), and Tabula Muris dataset. (B) Confusion matrix when using the learned classification tree to predict the labels of the Rosenberg dataset.

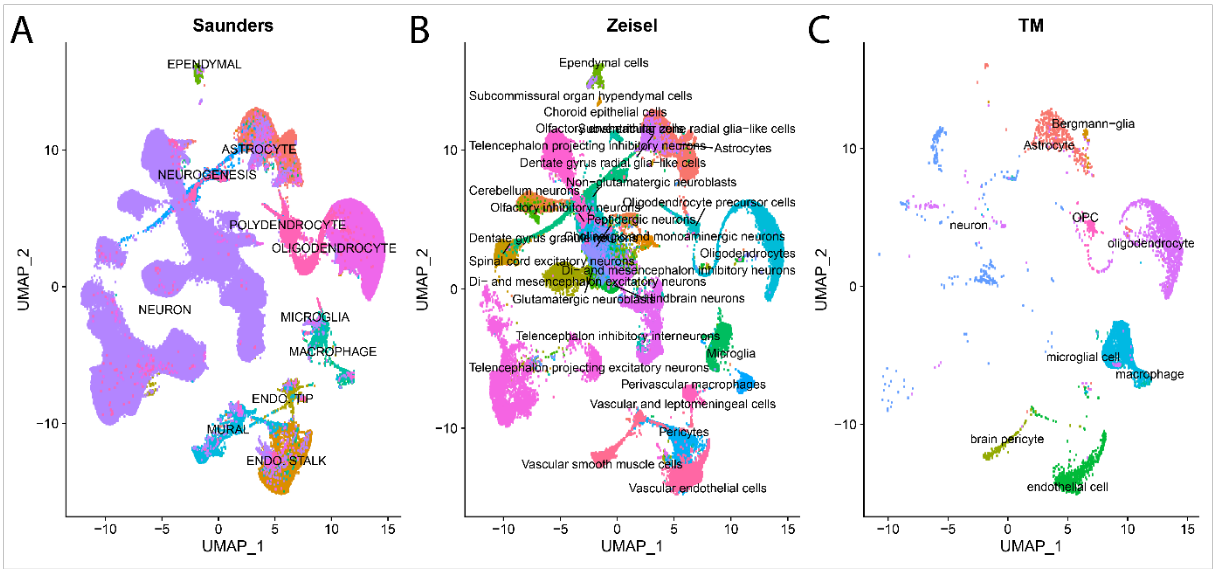


**Figure S21** UMAP embeddings of (A) Saunders, (B) Zeisel (high resolution), (C) Tabula Muris after data integration.

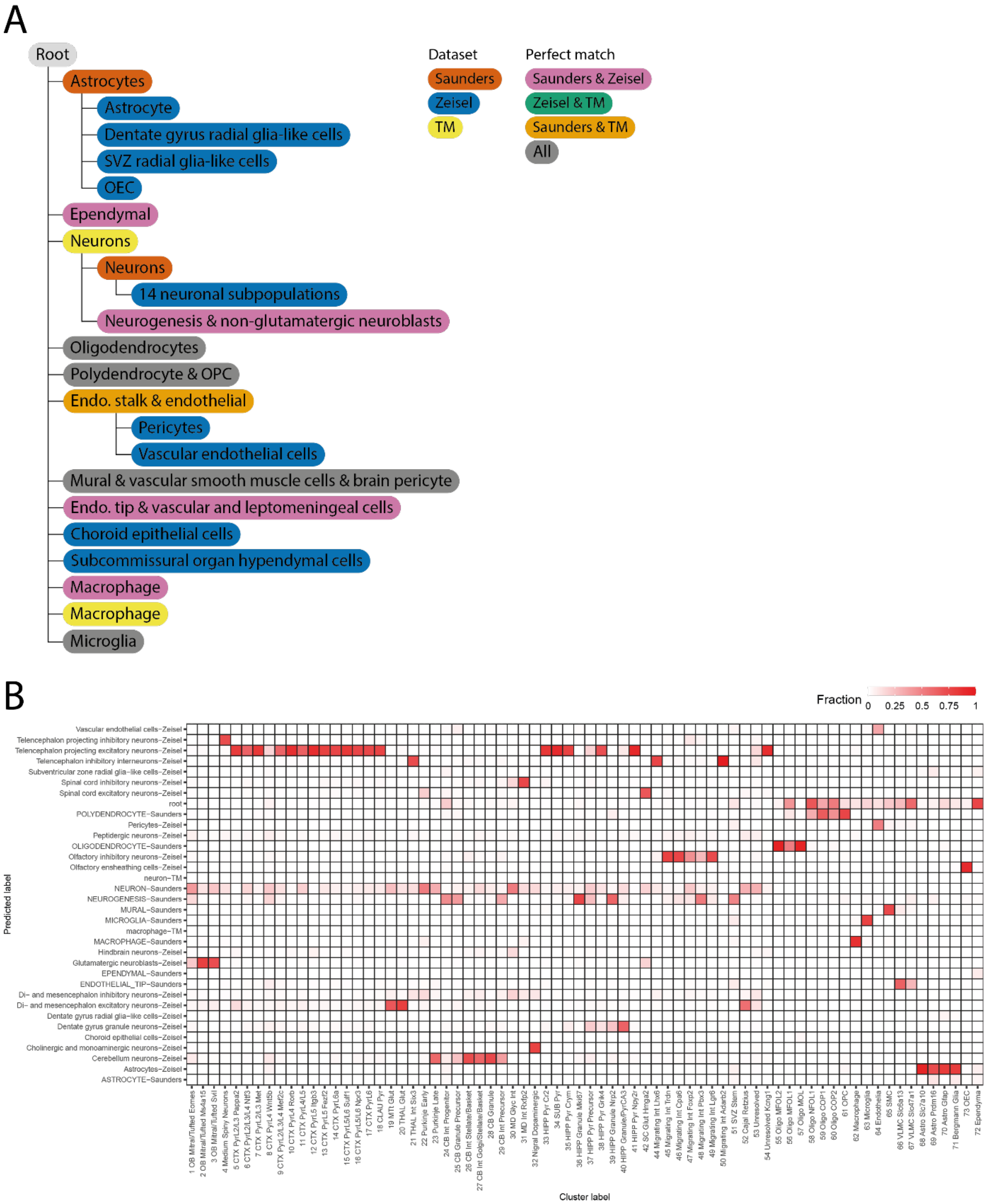


**Figure S22** (A) Learned hierarchy when applying *scHPL* with a linear SVM to the Saunders, Zeisel (high resolution), and Tabula Muris dataset. The 14 neuronal populations in the tree include: olfactory inhibitory neurons, cholinergic and monoaminergic neurons, glutamatergic neuroblasts, di- and mesencephalon inhibitory neurons, cerebellum neurons, peptidergic neurons, spinal cord inhibitory neurons, di- and mesencephalon excitatory neurons, spinal cord excitatory neurons, telencephalon projecting excitatory neurons, hindbrain neurons, dentate gyrus granule neurons, telencephalon inhibitory interneurons, and telencephalon projecting inhibitory neurons. Bergmann glia (TM) and astrocytes (TM) are missing from the tree. (B) Confusion matrix when using the learned classification tree to predict the labels of the Rosenberg dataset.

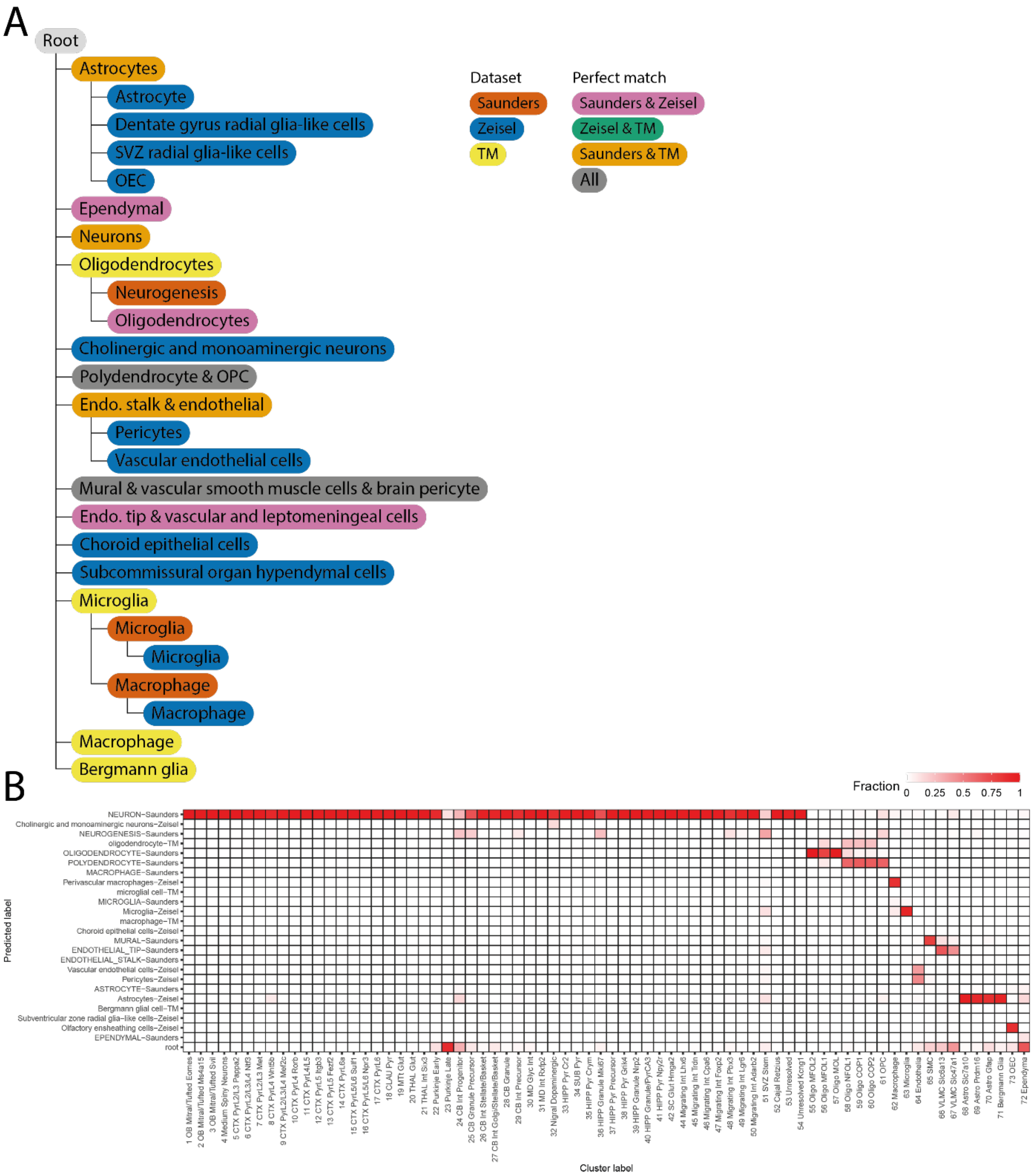


**Figure S23** Result one-class SVM on the high resolution. Almost all neuronal cell populations are missing. Fourteen neuronal populations are missing from the tree: olfactory inhibitory neurons, glutamatergic neuroblasts, non-glutamatergic neuroblasts, di- and mesencephalon inhibitory neurons, cerebellum neurons, peptidergic neurons, spinal cord inhibitory neurons, di- and mesencephalon excitatory neurons, spinal cord excitatory neurons, telencephalon projecting excitatory neurons, hindbrain neurons, dentate gyrus granule neurons, telencephalon inhibitory interneurons, and telencephalon projecting inhibitory neurons. (B) Confusion matrix when using the learned classification tree to predict the labels of the Rosenberg dataset.

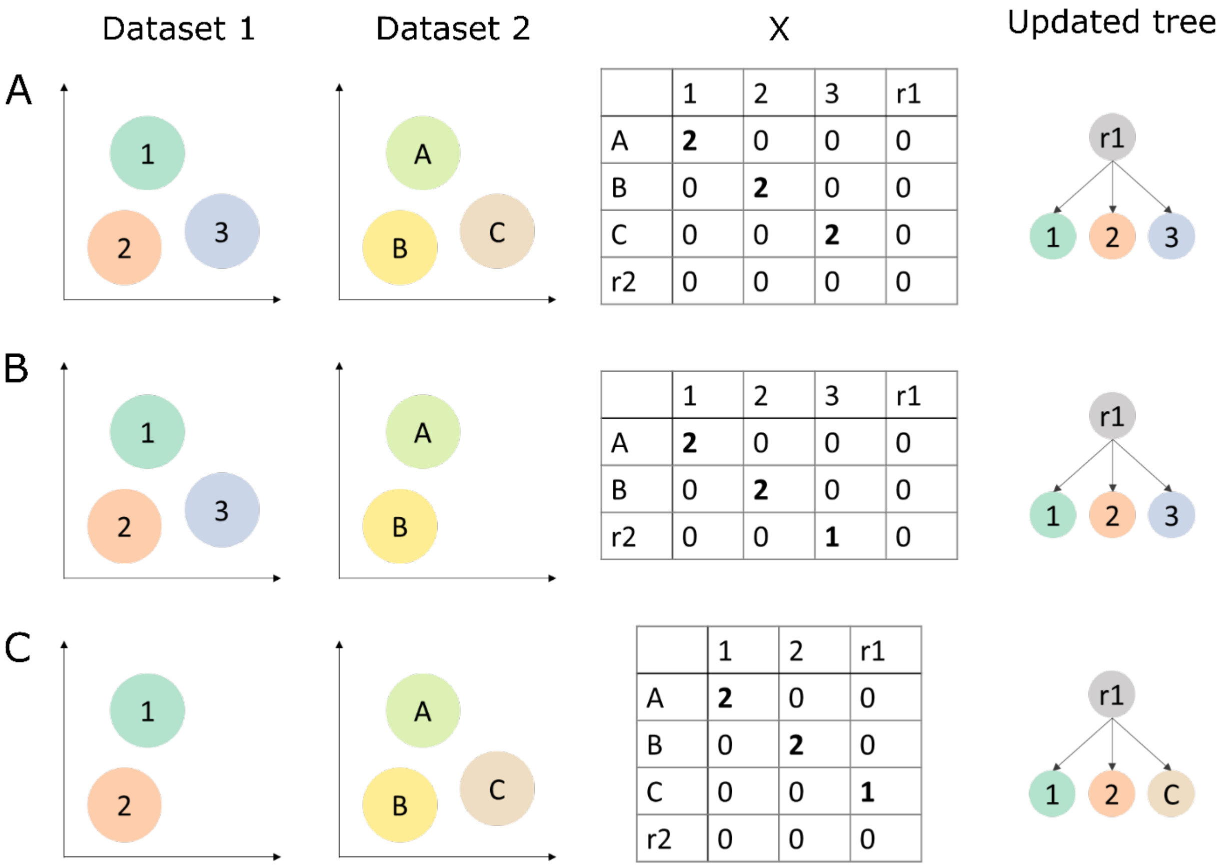


A

| | 1 | 2 | 3 | r1 |
|---|---|---|---|---|
| A | **2** | 0 | 0 | 0 |
| B | 0 | **2** | 0 | 0 |
| C | 0 | 0 | **2** | 0 |
| r2 | 0 | 0 | 0 | 0 |

B

| | 1 | 2 | 3 | r1 |
|---|---|---|---|---|
| A | **2** | 0 | 0 | 0 |
| B | 0 | **2** | 0 | 0 |
| r2 | 0 | 0 | **1** | 0 |

C

| | 1 | 2 | r1 |
|---|---|---|---|
| A | **2** | 0 | 0 |
| B | 0 | **2** | 0 |
| C | 0 | 0 | **1** |
| r2 | 0 | 0 | 0 |

**Figure S24** Schematic examples of the simple scenarios. For each scenario, we show what the cell populations in the two datasets could look like, X and the updated tree.

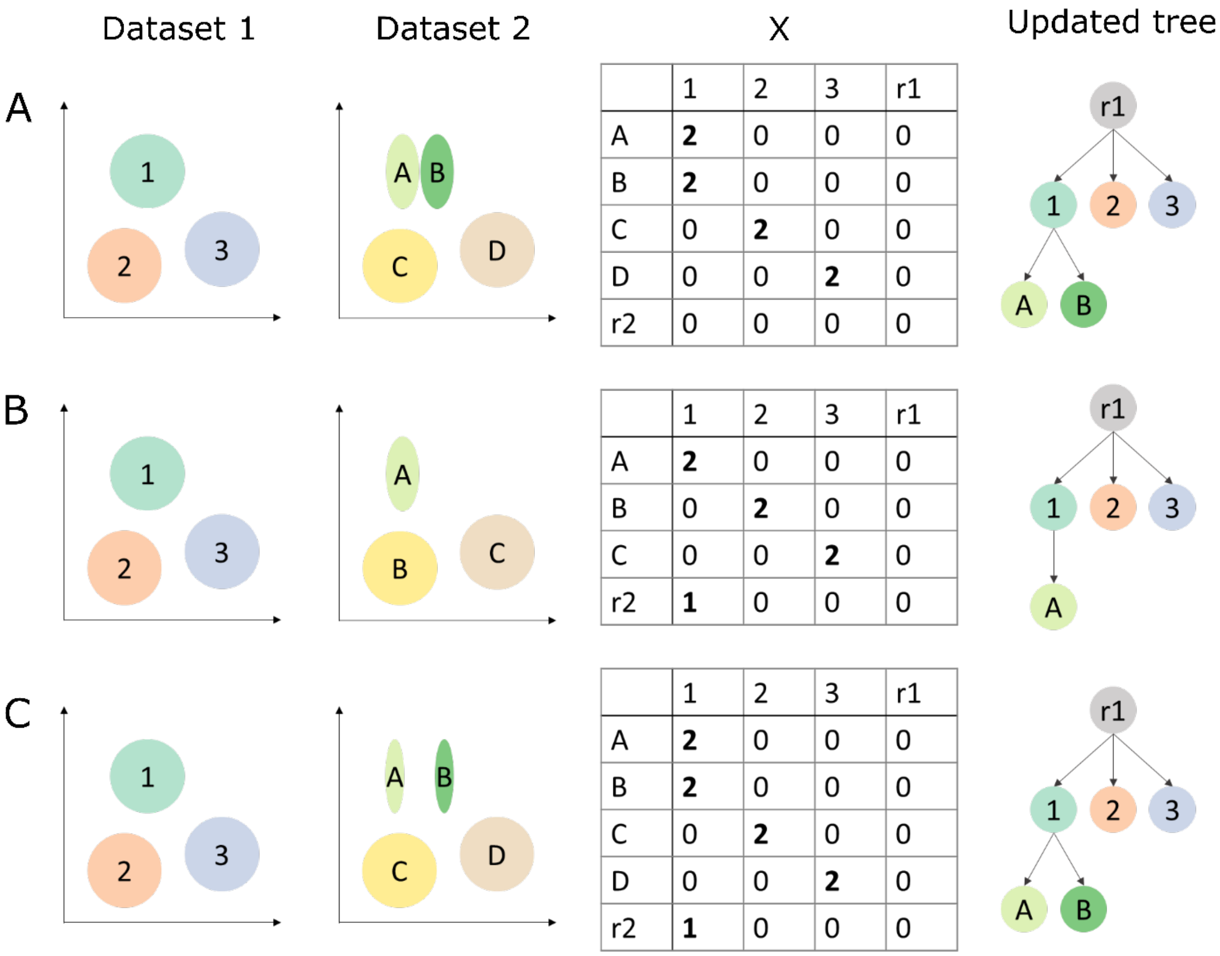


| | 1 | 2 | 3 | r1 |
|---|---|---|---|---|
| A | **2** | 0 | 0 | 0 |
| B | **2** | 0 | 0 | 0 |
| C | 0 | **2** | 0 | 0 |
| D | 0 | 0 | **2** | 0 |
| r2 | 0 | 0 | 0 | 0 |

| | 1 | 2 | 3 | r1 |
|---|---|---|---|---|
| A | **2** | 0 | 0 | 0 |
| B | 0 | **2** | 0 | 0 |
| C | 0 | 0 | **2** | 0 |
| r2 | **1** | 0 | 0 | 0 |

| | 1 | 2 | 3 | r1 |
|---|---|---|---|---|
| A | **2** | 0 | 0 | 0 |
| B | **2** | 0 | 0 | 0 |
| C | 0 | **2** | 0 | 0 |
| D | 0 | 0 | **2** | 0 |
| r2 | **1** | 0 | 0 | 0 |

**Figure S25** Schematic examples of the colsums scenarios. For each scenario, we show what the cell populations in the two datasets could look like, X and the updated tree.

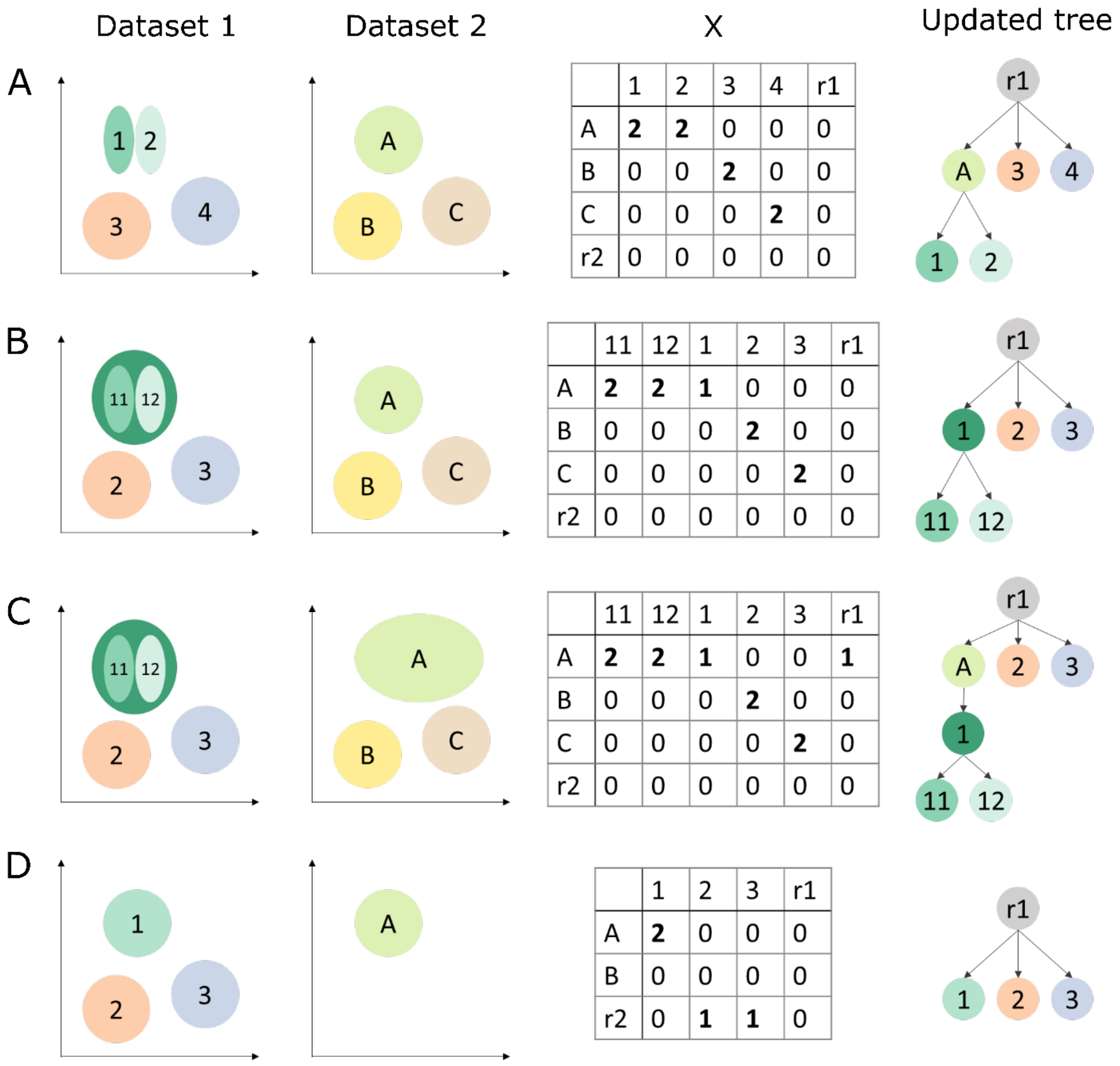


A

| | 1 | 2 | 3 | 4 | r1 |
|---|---|---|---|---|---|
| A | **2** | **2** | 0 | 0 | 0 |
| B | 0 | 0 | **2** | 0 | 0 |
| C | 0 | 0 | 0 | **2** | 0 |
| r2 | 0 | 0 | 0 | 0 | 0 |

B

| | 11 | 12 | 1 | 2 | 3 | r1 |
|---|---|---|---|---|---|---|
| A | **2** | **2** | **1** | 0 | 0 | 0 |
| B | 0 | 0 | 0 | **2** | 0 | 0 |
| C | 0 | 0 | 0 | 0 | **2** | 0 |
| r2 | 0 | 0 | 0 | 0 | 0 | 0 |

C

| | 11 | 12 | 1 | 2 | 3 | r1 |
|---|---|---|---|---|---|---|
| A | **2** | **2** | **1** | 0 | 0 | **1** |
| B | 0 | 0 | 0 | **2** | 0 | 0 |
| C | 0 | 0 | 0 | 0 | **2** | 0 |
| r2 | 0 | 0 | 0 | 0 | 0 | 0 |

D

| | 1 | 2 | 3 | r1 |
|---|---|---|---|---|
| A | **2** | 0 | 0 | 0 |
| B | 0 | 0 | 0 | 0 |
| r2 | 0 | **1** | **1** | 0 |

**Figure S26** Schematic examples of the rowsums scenarios. For each scenario, we show what the cell populations in the two datasets could look like, X and the updated tree.

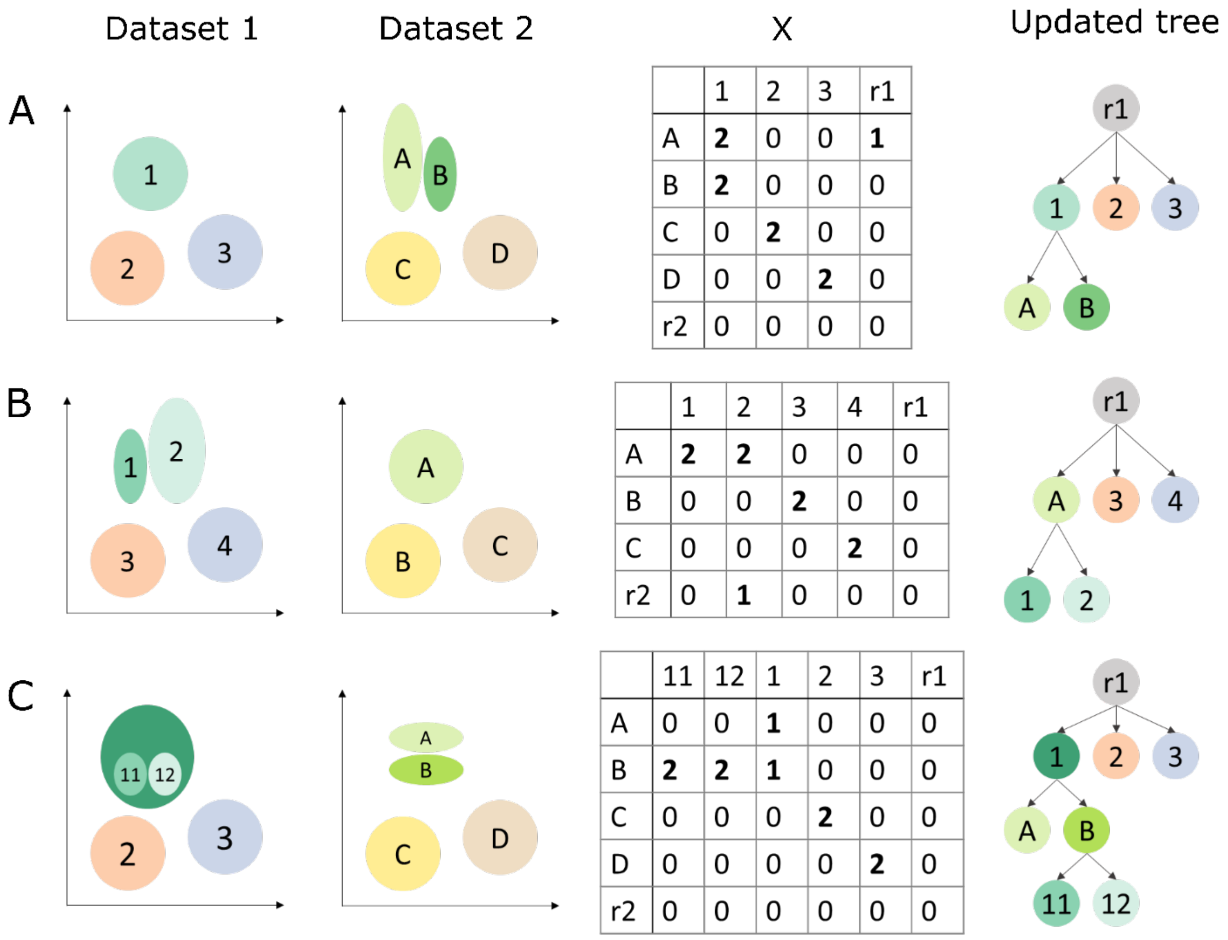


A:

| | 1 | 2 | 3 | r1 |
|---|---|---|---|---|
| A | **2** | 0 | 0 | **1** |
| B | **2** | 0 | 0 | 0 |
| C | 0 | **2** | 0 | 0 |
| D | 0 | 0 | **2** | 0 |
| r2 | 0 | 0 | 0 | 0 |

B:

| | 1 | 2 | 3 | 4 | r1 |
|---|---|---|---|---|---|
| A | **2** | **2** | 0 | 0 | 0 |
| B | 0 | 0 | **2** | 0 | 0 |
| C | 0 | 0 | 0 | **2** | 0 |
| r2 | 0 | **1** | 0 | 0 | 0 |

C:

| | 11 | 12 | 1 | 2 | 3 | r1 |
|---|---|---|---|---|---|---|
| A | 0 | 0 | **1** | 0 | 0 | 0 |
| B | **2** | **2** | **1** | 0 | 0 | 0 |
| C | 0 | 0 | 0 | **2** | 0 | 0 |
| D | 0 | 0 | 0 | 0 | **2** | 0 |
| r2 | 0 | 0 | 0 | 0 | 0 | 0 |

**Figure S27** Schematic examples of the complex scenarios. For each scenario, we show what the cell populations in the two datasets could look like, X and the updated tree.

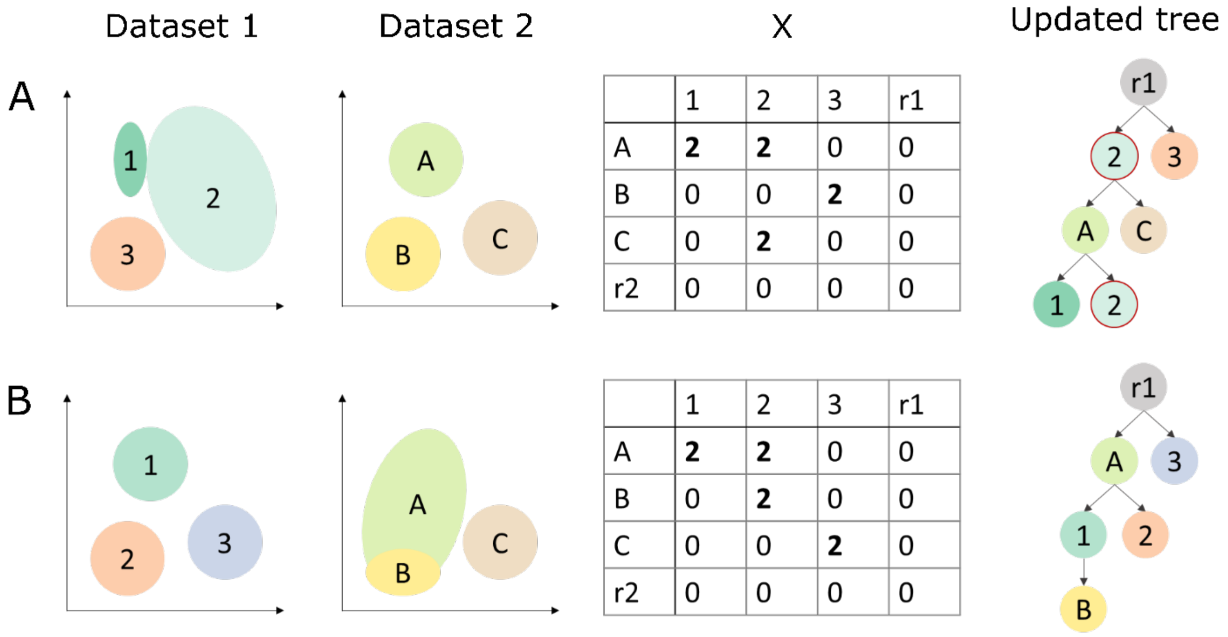


A:

| | 1 | 2 | 3 | r1 |
|---|---|---|---|---|
| A | **2** | **2** | 0 | 0 |
| B | 0 | 0 | **2** | 0 |
| C | 0 | **2** | 0 | 0 |
| r2 | 0 | 0 | 0 | 0 |

B:

| | 1 | 2 | 3 | r1 |
|---|---|---|---|---|
| A | **2** | **2** | 0 | 0 |
| B | 0 | **2** | 0 | 0 |
| C | 0 | 0 | **2** | 0 |
| r2 | 0 | 0 | 0 | 0 |

**Figure S28** Schematic examples of the impossible scenarios. For each scenario, we show what the cell populations in the two datasets could look like, X and why the updated tree is not possible.

**Supplementary tables**

**Table S1** Explanation of the most common matching scenarios.

| Scenario | Biological explanation | Identification in the matching matrix (*X*) | Solution |
|---|---|---|---|
| Perfect match | Population *i* of dataset 1 exactly matches population *j* of dataset 2 | $X_{j,i} = 1$ or *2* rest of row *j* and column *i* is zero | Rename labels of population *j* to population *i* |
| Splitting populations | Multiple populations from dataset 2 are subpopulations of population *i* of dataset 1 | Multiple non-zero values in column *i* of *X* | Add corresponding populations from dataset 2 as children to population *i* |
| Merging populations | Multiple populations from dataset 1 are subpopulations of population *j* of dataset 2 | Multiple non-zero values in column *j* of *X* | Add population *j* as a parent to the corresponding populations of dataset 1 |
| New population | Dataset 2 contains a new, unseen population *j* | $X_{j,root1} = 1$, the rest of row *j* is zero | Add population *j* as a child to the root node. |

**Table S2** Labels of the simulated dataset when testing tree construction

| Original label | Label Batch 1 | Label Batch 2 | Label Batch 3 |
|---|---|---|---|
| Group1 | Group12 | Group1 | Group1 |
| Group2 | Group12 | Group2 | Group2 |
| Group3 | Group3 | Group3 | Group3 |
| Group4 | Group456 | Group4 | Group4 |
| Group5 | Group456 | Group56 | Group5 |
| Group6 | Group456 | Group56 | Group6 |

**Table S3** Labels of PBMC-FACS dataset when testing tree construction

| Original label | Label Batch 1 | Label Batch 2 | Label Batch 3 |
|---|---|---|---|
| CD14+ Monocytes | CD14+ Monocytes | CD14+ Monocytes | CD14+ Monocytes |
| CD19+ B-cells | CD19+ B-cells | CD19+ B-cells | CD19+ B-cells |
| CD56+ NK cells | CD56+ NK cells | CD56+ NK cells | CD56+ NK cells |
| CD4+ T-cells | T-cells | CD4+ T-cells | - |
| CD4+/CD25+ reg. T-cells | T-cells | - | CD4+/CD25+ reg. T-cells |
| CD4+/CD45RA+/CD25- naive T-cells | T-cells | - | CD4+/CD45RA+/CD25- naive T-cells |
| CD4+/CD45RO+ mem. T-cells | T-cells | - | CD4+/CD45RO+ mem. T-cells |
| CD8+ T-cells | T-cells | CD8+ T-cells | - |
| CD8+/CD45RA+ naïve T-cells | T-cells | - | CD8+/CD45RA+ naïve T-cells |

**Table S4** Confusion matrix of the linear SVM on the PBMC data. Here, the linear SVM was trained using the predefined hematopoietic tree.

| | CD14+ MC | CD19+ B | CD34+ | CD56+ NK | CD4+ T | CD4+ T Reg | CD4+ T Naive | CD4+ T Memory | CD8+ T | CD8+ T Naive | root |
|---|---|---|---|---|---|---|---|---|---|---|---|
| **CD14+ MC** | 1957 | 1 | 0 | 1 | 2 | 15 | 0 | 1 | 0 | 0 | 23 |
| **CD19+ B** | 0 | 1998 | 0 | 0 | 0 | 0 | 0 | 1 | 0 | 0 | 1 |
| **CD34+** | 1 | 26 | 1794 | 1 | 0 | 0 | 0 | 0 | 6 | 0 | 172 |
| **CD56+ NK** | 0 | 0 | 2 | 1988 | 0 | 0 | 0 | 1 | 6 | 0 | 3 |
| **CD4+ T** | 1 | 0 | 0 | 0 | 191 | 1010 | 790 | 3 | 0 | 2 | 3 |
| **CD4+ T Reg** | 0 | 0 | 0 | 0 | 113 | 1704 | 175 | 4 | 0 | 0 | 4 |
| **CD4+ T Naive** | 1 | 1 | 0 | 0 | 94 | 140 | 1751 | 1 | 1 | 9 | 2 |
| **CD4+ T Memory** | 0 | 0 | 3 | 0 | 1 | 33 | 14 | 1888 | 32 | 26 | 3 |
| **CD8+ T** | 0 | 0 | 0 | 0 | 1 | 5 | 0 | 44 | 1884 | 65 | 1 |
| **CD8+ T Naive** | 0 | 1 | 0 | 0 | 3 | 0 | 13 | 36 | 31 | 1916 | 0 |

**Table S5** Confusion matrix of the linear SVM on the PBMC data. Here, the linear SVM was trained on the learned tree. The CD4+ memory T-cells are a subpopulation of CD8+ T-cells now.

| | CD14+ MC | CD19+ B | CD34+ | CD56+ NK | CD4+ T | CD4+ T Reg | CD4+ T Naive | CD4+ T Memory | CD8+ T | CD8+ T Naive | root |
|---|---|---|---|---|---|---|---|---|---|---|---|
| **CD14+ MC** | 1957 | 1 | 0 | 1 | 0 | 17 | 0 | 1 | 0 | 0 | 23 |
| **CD19+ B** | 0 | 1998 | 0 | 0 | 0 | 0 | 0 | 0 | 0 | 1 | 1 |
| **CD34+** | 1 | 26 | 1794 | 1 | 0 | 0 | 0 | 2 | 0 | 4 | 172 |
| **CD56+ NK** | 0 | 0 | 2 | 1988 | 0 | 0 | 0 | 7 | 0 | 0 | 3 |
| **CD4+ T** | 1 | 0 | 0 | 0 | 0 | 1116 | 875 | 5 | 0 | 0 | 3 |
| **CD4+ T Reg** | 0 | 0 | 0 | 0 | 0 | 1794 | 200 | 2 | 0 | 0 | 4 |
| **CD4+ T Naive** | 1 | 1 | 0 | 0 | 0 | 133 | 1860 | 2 | 0 | 1 | 2 |
| **CD4+ T Memory** | 0 | 0 | 3 | 0 | 0 | 4 | 0 | 1972 | 0 | 18 | 3 |
| **CD8+ T** | 0 | 0 | 0 | 0 | 0 | 2 | 0 | 774 | 0 | 1223 | 1 |
| **CD8+ T Naive** | 0 | 1 | 0 | 0 | 0 | 0 | 3 | 24 | 0 | 1972 | 0 |

**Table S6** Labels of the simulated dataset when testing tree construction with missing cell populations

| Original label | Label Batch 1 | Label Batch 2 | Label Batch 3 |
|---|---|---|---|
| Group1 | Group12 | Group1 | Group1 |
| Group2 | Group12 | Group2 | Group2 |
| Group3 | Group3 | Group3 | Group3 |
| Group4 | Group456 | Group4 | Group4 |
| Group5 | Group456 | - | Group5 |
| Group6 | Group456 | Group6 | Group6 |

**Table S7** Characteristics of the brain datasets

| | Year | Protocol | Number of cells | Number of cell populations |
|---|---|---|---|---|
| **Tabula Muris** | 2018 | Smart-seq2 | 7,856 | 9 |
| **Rosenberg** | 2018 | SPLiT-seq | 76,322 | 73 |
| **Zeisel** | 2018 | 10Xv1 | 85,621 | 11, 30 |
| **Saunders** | 2018 | Drop-seq | 389,439 | 11, 437 |
